# Incomplete bunyavirus particles contribute to within-host spread and between-host transmission

**DOI:** 10.1101/2022.03.07.483181

**Authors:** Erick Bermúdez-Méndez, Kirsten F. Bronsvoort, Mark P. Zwart, Sandra van de Water, Ingrid Cárdenas-Rey, Rianka P. M. Vloet, Constantianus J. M. Koenraadt, Gorben P. Pijlman, Jeroen Kortekaas, Paul J. Wichgers Schreur

## Abstract

Bunyaviruses lack a specific mechanism to ensure the incorporation of a complete set of genome segments into each virion, explaining the generation of incomplete virus particles lacking one or more genome segments. Such incomplete virus particles, which may represent the majority of particles produced, are generally considered to interfere with virus infection and spread. Using the three-segmented Rift Valley fever virus as a model bunyavirus, we here show that two distinct incomplete virus particle populations that are unable to spread autonomously, are able to efficiently complement each other in both mammalian and insect cells following co-infection. We further show that incomplete virus particles are capable of co-infecting mosquitoes, resulting in the rescue of infectious virus that is able to disseminate to the mosquito salivary glands. Our findings reveal a significant role of incomplete particles in within-host spread and between-host transmission, reminiscent of the life cycle of multipartite viruses.

## Introduction

Segmented and multipartite viruses have genomes divided over multiple segments. The classical paradigm in virology states that segmented viruses package all their genome segments into a single virus particle, whereas multipartite viruses package each genome segment into a distinct virus particle^1^. To ensure a productive infection, it is generally accepted that multipartite viruses (mainly found to infect plants and fungi) rely on co-infection of the same cell with a set of complementing particles, each particle containing a different genome segment^1,2^. Alternatively, complementation can occur at the tissue level, as proposed in a recent study with the plant-infecting faba bean necrotic stunt virus (FBNSV, family *Nanoviridae*). FBNSV was shown to complement its missing genome segments by export and distribution of viral mRNAs and proteins across interconnected neighboring cells^3^. By contrast, it has been thought that segmented viruses (mainly found to infect animals) solely rely on individual cells as units of viral replication, and thus have to carry at least one copy of each genome segment within a single virus particle to ensure the delivery of a complete genome^4,5^.

Under this traditional view on segmented viruses, it seems reasonable to expect a selective genome packaging strategy that facilitates the generation of progeny virus particles containing a complete set of genome segments. Otherwise, in the absence of such a selective packaging strategy, only a minor fraction of the particles being produced would contain a complete set of genome segments. The production of particles lacking one or more viral genome segments can compromise efficient viral spread, as they are incapable of initiating a productive infection on their own. Influenza A virus (IAV, family *Orthomyxoviridae*) is the prime example of a segmented virus having a highly selective genome packaging mechanism, in which intersegment interactions facilitate the assembly of its eight-segmented genome into a supramolecular complex that is incorporated inside new virus particles^6–12^. However, not all segmented viruses seem to employ such an orchestrated genome packaging process, leaving a gap in our understanding of the full replication cycle of, for instance, viruses belonging to the order *Bunyavirales*.

Bunyaviruses are enveloped, negative- or ambi-sense, single-stranded RNA viruses with a genome divided over two to six segments. These viruses are transmitted by arthropods or rodents, and can infect a wide variety of hosts, including mammals, birds, reptiles and plants^13^. Recently, using a combined single-molecule fluorescence *in situ* hybridization (smFISH)-immunofluorescence approach, we showed that the genome packaging processes of two members of the *Bunyavirales*, Rift Valley fever virus (RVFV, family *Phenuiviridae*, genus *Phlebovirus*) and Schmallenberg virus (SBV, family *Peribunyaviridae*, genus *Orthobunyavirus*), are not tightly controlled. Such non-selective packaging mechanism results in mixed virus progeny populations that consist of a minor fraction (below 25%) of complete particles (*i*.*e*. containing a complete set of all three genome segments: S, M and L) and a large fraction (above 75%) of empty and incomplete particles (*i*.*e*. lacking one or more genome segments)^14,15^. Despite the apparent inefficient genome packaging of RVFV and SBV, these viruses are able to spread efficiently within and between their mammalian and arthropod hosts. Interestingly, the spread of RVFV and SBV seems to be unaffected by the non-selective packaging strategy and/or is compensated by benefits to viral replication or spread through yet unknown mechanisms. Due to the fact that bunyaviruses only generate a small fraction of complete particles, we hypothesize that particles with an incomplete set of genome segments may still contribute to efficient bunyavirus spread by genetic complementation after co-infecting the same cell.

In this study, we used RVFV variants encoding fluorescent reporter proteins to investigate cell susceptibility to simultaneous infection by more than one virus particle. Subsequently, we assessed whether within-host genome complementation could occur in mammalian and insect cells by generating different two-segmented incomplete virus particle populations that depend entirely on co-infection for the production of progeny virions. We further investigated whether particles with an incomplete set of genome segments can complement each other *in vivo* in the mosquito vector. Lastly, we mathematically modeled diverse infection scenarios to estimate under what conditions incomplete particles substantially contribute to virus spread. The results of our study point towards a previously overlooked, yet significant role of incomplete particles in the bunyavirus life cycle, showing that incomplete virus particles can contribute to both within-host spread and between-host transmission.

## Results

### Efficient co-infection of three- and two-segmented RVFV reporter viruses in both mammalian and insect cells

In vertebrates, genetic complementation of virus particles with a distinct genome composition strictly relies on their ability to co-infect the same cell. To assess if mammalian and insect cells are susceptible to RVFV co-infection, we generated two recombinant three-segmented RVFV variants encoding either eGFP or mCherry2 in place of the NSs gene, using a T7 polymerase-based reverse genetics system (**Fig. 1a**). Cells infected with the individual reporter viruses showed abundant expression of the respective fluorescent protein (**Fig. 1b**), providing a suitable strategy to track virus infection and identify co-infected cells through the detection of co-localized fluorescent signals. Moreover, RVFV-eGFP and RVFV-mCherry2 replicated with almost identical growth kinetics, reaching high titers already 24 h post-infection and peaking at 48 h post-infection (**Fig. 1c**). Upon simultaneous inoculation with RVFV-eGFP and RVFV-mCherry2, both BSR-T7/5 (hamster) and C6/36 (mosquito) cells were found to be susceptible to co-infection, as clearly evidenced by the co-localization of green and red fluorescent signal in a fraction of the cell population (**Fig. 1d-e**).

**Fig. 1.**
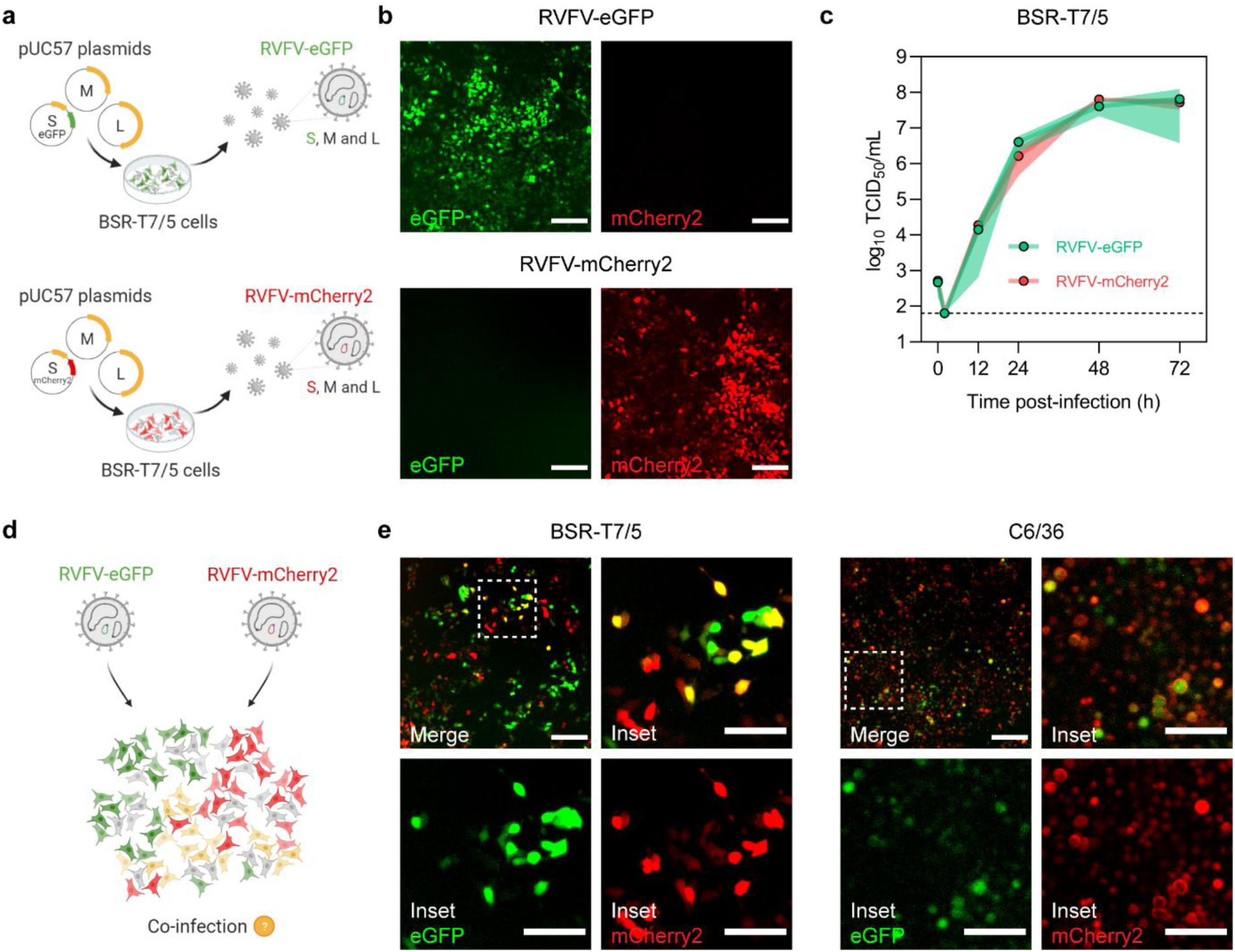
Co-infection of mammalian and insect cells with recombinant three-segmented RVFV reporter viruses. **a** Schematic representation of the T7 polymerase-based reverse genetics system. Fluorescently marked variants of RVFV were generated by simultaneous transfection of BSR-T7/5 cells with the transcription plasmids pUC57_S, pUC57_M and pUC57_L encoding the RVFV-35/74 S, M and L genome segments, respectively, in antigenomic-sense orientation. The pUC57_S plasmid additionally encoded for either eGFP or mCherry2 in place of the NSs gene. **b** Direct fluorescence detection of BSR-T7/5 cells infected with either RVFV-eGFP (green) or RVFV-mCherry2 (red) (MOI 0.5) at 24 h post-infection. **c** Growth kinetics of RVFV-eGFP and RVFV-mCherry2 after infection of BSR-T7/5 cells at an MOI of 0.01. Virus titers were determined with an end-point dilution assay (fluorescence microscopy read-out). Dots represent means of biological replicates (n = 3) at each time point and the shaded area represents the standard deviation. The dashed line indicates the limit of detection (10^1.8^ TCID_50_/mL). **d** Schematic representation of the simultaneous infection of mammalian (BSR-T7/5) or insect (C6/36) cells with RVFV-eGFP and RVFV-mCherry2. **e** Direct fluorescence detection of BSR-T7/5 and C6/36 cells co-infected with RVFV-eGFP and RVFV-mCherry2 (MOI 0.5 for each virus) at 24 h (BSR-T7/5) or 72 h (C6/36) post-infection. Inset images are magnifications of a region of interest (indicated as a dashed box). Co-infected cells co-express eGFP (green) and mCherry2 (red) and thus appear yellow. Scale bars, 200 μm (inset images 100 μm).

Simultaneous infections with RVFV-eGFP and RVFV-mCherry2 allowed us to qualitatively confirm cell susceptibility to co-infection (**Fig. 1e**). However, the fast spread of three-segmented RVFV over multiple infection cycles impeded us to assess accurately how often these co-infection events are actually taking place. To overcome this, we again used the T7 polymerase-based reverse genetics system to generate two non-spreading incomplete RVFV particle populations lacking the M segment and encoding a fluorescent protein (FP), either eGFP or mCherry2, in the S segment in place of the NSs gene (iRVFV-SL-FP) (**Fig. 2a**). The generation of iRVFV-SL-eGFP has been previously reported by our group (initially termed RRPs)^16^, but the generation of iRVFV-SL-mCherry2 is first reported as part of this work. Both particle populations are produced following complementation with an expression plasmid encoding the structural glycoproteins (Gn and Gc) normally encoded by the M segment. Despite the absence of the M segment, the S and L genome segments encoding for the nucleocapsid (N) protein and the RNA-dependent RNA-polymerase (RdRp or L protein), respectively, are sufficient to support replication and transcription of the S and L segments upon infection of naive cells. Importantly, because the M segment encoding for the Gn and Gc glycoproteins is absent in these naive cells, infection with iRVFV-SL particles does not lead to assembly and release of progeny virus. Direct fluorescence microscopy combined with an immunofluorescence assay to detect Gn clearly showed that cells infected with iRVFV-SL-eGFP or iRVFV-SL-mCherry2 have abundant expression of the respective fluorescent protein but no expression of Gn (**Fig. 2b**). Furthermore, passaging the supernatant of cells infected with iRVFV-SL-FP to naive cells did not result in the expression of eGFP, mCherry2, or Gn, confirming that these particles are incomplete and not able to spread due to the lack of the M genome segment (**Fig. 2c**).

**Fig. 2.**
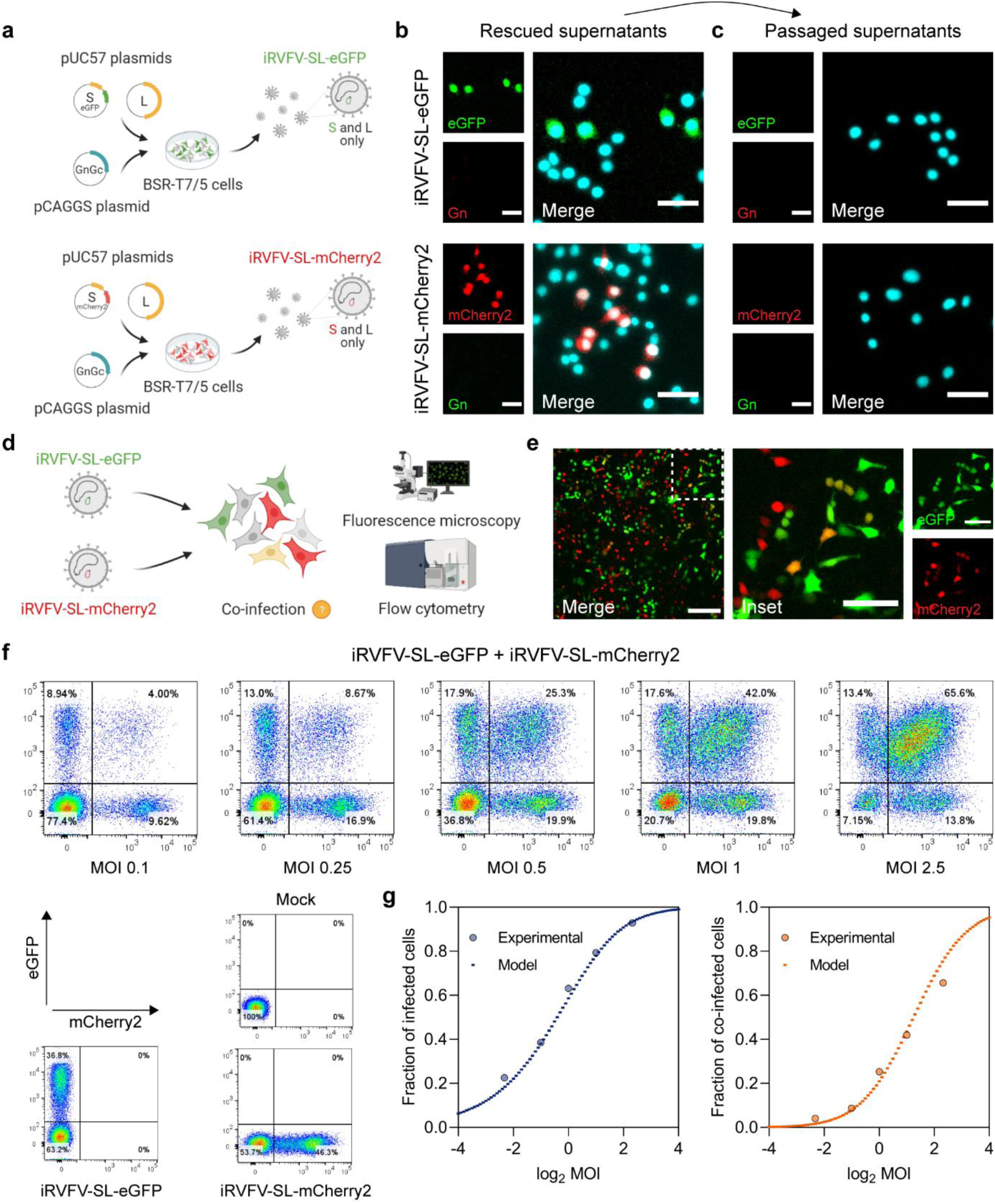
Quantitative assessment of mammalian cells co-infected with non-spreading two-segmented RVFV reporter viruses. **a** Schematic representation of the reverse genetics system used to create two-segmented RVFV reporter viruses. Incomplete RVFV-SL particles were generated by co-transfection of BSR-T7/5 cells with the transcription plasmids pUC57_S, encoding either eGFP or mCherry2 in place of the NSs gene, pUC57_L and the protein expression plasmid pCAGGS_NSmGnGc. **b** Immunofluorescence assay for detection of BSR-T7/5 cells infected with iRVFV-SL-eGFP or iRVFV-SL-mCherry2 (MOI 0.5) at 24 h post-infection. **c** Supernatants from cells primarily infected with the incomplete RVFV particles in **b** were passaged onto naive BSR-T7/5 cells. Cells were examined at 24 h post-infection for expression of eGFP (green) and mCherry2 (red) by fluorescence microscopy. Expression of the Gn glycoprotein (green or red, depending on the virus) was detected with rabbit polyclonal anti-Gn serum in combination with FITC-conjugated (green) or Alexa Fluor 568-conjugated (red) secondary antibodies. Cell nuclei (cyan) were visualized with DAPI. Scale bars, 50 μm. **d** Schematic representation of the simultaneous infection of mammalian (BSR-T7/5) cells with non-spreading iRVFV-SL-eGFP and iRVFV-SL-mCherry2 particles. Co-infections were done at increasing MOI (ranging from 0.1 to 2.5 for each virus). After 48 h, cells were examined by fluorescence microscopy and fixed for flow cytometry analysis. **e** Direct fluorescence detection of BSR-T7/5 cells co-infected (MOI of 0.5 for each virus) with the two-segmented incomplete particle populations. The inset image is a magnification of a region of interest (indicated as a dashed box). Co-infected cells co-express eGFP (green) and mCherry2 (red) and thus appear yellow. Scale bars, 200 μm (inset images 100 μm). **f** Cells expressing eGFP, mCherry2 or both were quantified by flow cytometry. Mock-infected cells and cells infected exclusively with only one population of incomplete particles (MOI 0.5) were used as controls. **g** Relationship between the fraction of infected (left) and co-infected (right) cells as a function of the MOI. Dots represent experimental data points. Dashed lines represent the predictions of a model based on the assumptions that genome segments are randomly packaged into virus particles and that host susceptibility is heterogeneous.

To quantify to what extent cells can be co-infected with the two different non-spreading fluorescent virus variants, we simultaneously infected BSR-T7/5 cells with iRVFV-SL-eGFP and iRVFV-SL-mCherry2 at increasing multiplicities of infection (MOIs, ranging from 0.1 to 2.5) (**Fig. 2d**). Through direct detection of co-localized eGFP and mCherry2 expression, we confirmed that these particles also could co-infect BSR-T7/5 cells (**Fig. 2e**). Next, using flow cytometry, we quantified the fraction of non-infected cells, singly-infected cells and co-infected cells (**Fig. 2f**). Mock-infected cells and cells infected with only one population of incomplete particles were the basis to gate the flow cytometry data. A clear double (eGFP and mCherry2)-positive cell population was identified after co-infection with both populations of incomplete particles. Interestingly, at each MOI tested, the percentage of infected and co-infected cells closely resembled that of a predictive mathematical model based on the assumptions that genome segments are randomly packaged into virus particles and that host susceptibility is heterogeneous (**Fig. 2g**). Moreover, the population of co-infected cells rose sharply with increasing MOI in a dose-response fashion, suggesting that there is no apparent mechanism leading to the exclusion of multiple particles entering the same cell in the present experimental setup.

### Successful generation of incomplete RVFV particles lacking the S genome segment

To study the potential role of incomplete particles in the bunyavirus life cycle, we needed at least two distinct populations of RVFV particles having incomplete but complementing genomes. Besides generating iRVFV-SL-FP particles, we also generated incomplete RVFV particles lacking the S segment (iRVFV-ML), by following a similar T7 polymerase-based reverse genetics strategy (**Fig. 3a**). Complete M and L genome segments were encoded in antigenomic-sense orientation by transcription plasmids, and the N protein was provided by an expression plasmid. Due to the absence of the S segment in the rescued particles, and consequently, the unavailability of N protein, infections with iRVFV-ML particles did not result in any appreciable viral genome replication. Accordingly, neither N nor Gn was detected in an immunofluorescence assay with cells infected with iRVFV-ML (**Fig. 3b**). Due to the non-replicating nature of RVFV particles lacking the S genome segment, the infectious titer of iRVFV-ML stocks could not be determined with a conventional virus titration assay. Therefore, we confirmed the genomic composition of rescued iRVFV-ML particles through quantification of the S, M and L genome segments via RT-qPCR (**Fig. 3c**). In both batches of rescued iRVFV-ML particles, high copy numbers of only the M and L genome segments were detected, whereas the S segment was not detected. Similarly, in samples containing iRVFV-SL-eGFP particles, only the S and L genome segments were present at high copy numbers and the M segment was not detected. All three genome segments (S, M and L) were detected in samples containing the three-segmented RVFV-eGFP used as control.

**Fig. 3.**
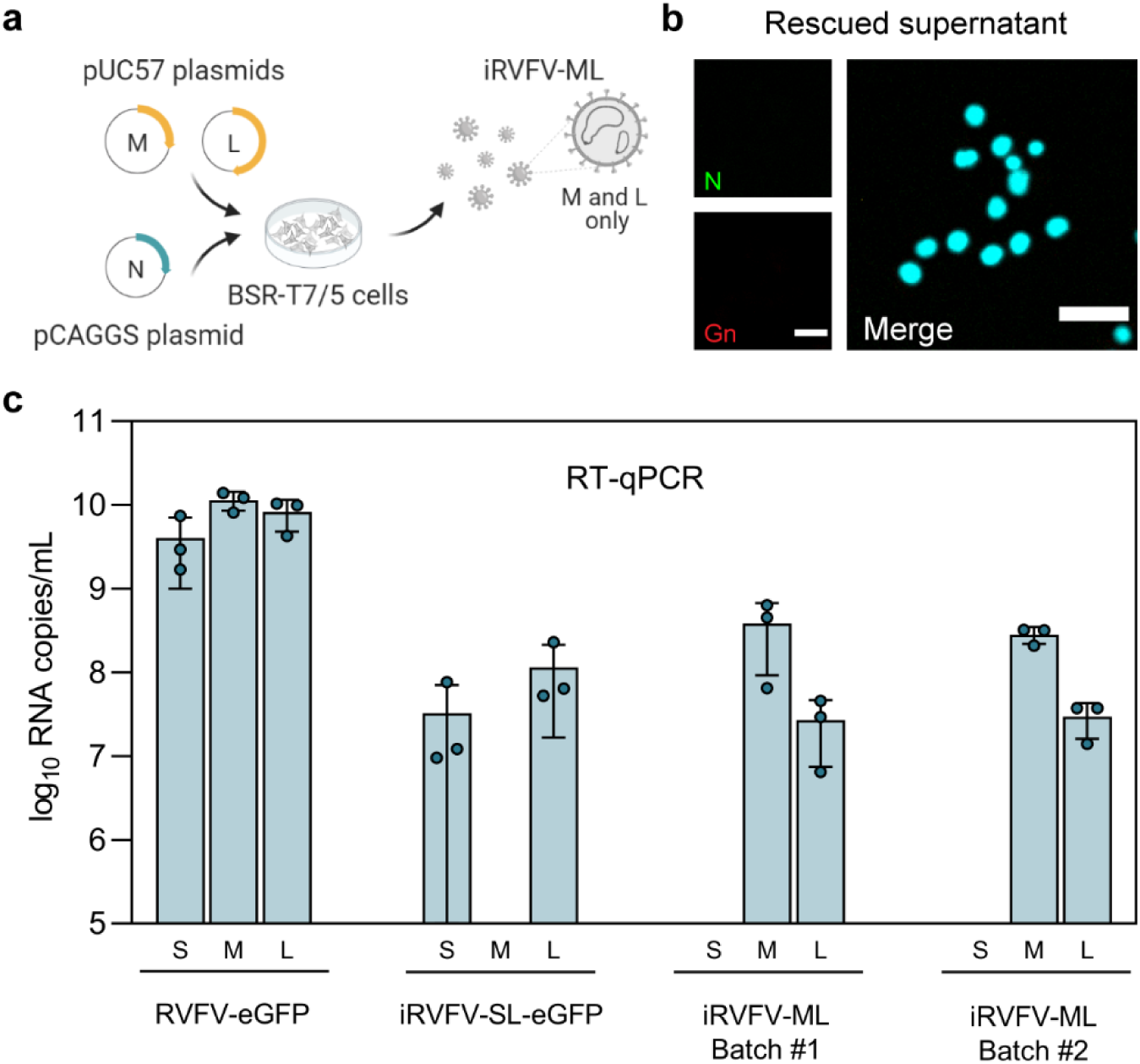
Generation of incomplete RVFV particles lacking the S genome segment. **a** Schematic representation of the reverse genetics system used to create incomplete RVFV-ML particles. iRVFV-ML particles were generated by co-transfection of BSR-T7/5 cells with the transcription plasmids pUC57_M, pUC57_L and the protein expression plasmid pCAGGS_N. **b** Immunofluorescence assay for detection of BSR-T7/5 cells exposed to iRVFV-ML (MOI 0.5) at 24 h post-infection. Expression of the N protein (green) was detected with a mouse anti-N monoclonal hybridoma in combination with Alexa Fluor Plus 488-conjugated secondary antibodies. Expression of the Gn glycoprotein (red) was detected with rabbit polyclonal anti-Gn serum in combination with Alexa Fluor 568-conjugated secondary antibodies. Cell nuclei (cyan) were visualized with DAPI. Scale bars, 50 μm. **c** Segment-specific RT-qPCR quantification of the genomic composition of two batches of rescued iRVFV-ML particles. Stock preparations of RVFV-eGFP and iRVFV-SL-eGFP were included as reference for comparison. Bars show means with SD. Dots represent replicates (n = 3 samples).

### Co-infection with complementing incomplete RVFV particles results in efficient virus replication and spread among mammalian and insect cells

Having generated two distinct populations of two-segmented RVFV particles with an incomplete but complementing genome, we next investigated whether exposing cells to a mixture of these two populations would result in genome complementation (SL + ML = SML) and subsequent rescue of three-segmented (SML) infectious virus (**Fig. 4a**). Rescue of infectious virus following incubation of cells exclusively with virus populations with an incomplete set of genome segments would strongly suggest a role for incomplete particles in virus replication and spread. To this end, BSR-T7/5 and C6/36 cells were simultaneously inoculated with iRVFV-SL-eGFP and iRVFV-ML at different MOIs (ranging from 0.001 to 0.25 for each virus). As shown previously in **Fig. 2b** and **Fig. 3b**, incubation of cells with only iRVFV-SL-eGFP or iRVFV-ML particles did not result in productive infections. Inoculation with iRVFV-SL-eGFP particles resulted in the expression of eGFP in single cells without spread to neighboring cells, and inoculation with iRVFV-ML particles did not result in the production of viral proteins. Remarkably, upon simultaneous inoculation with iRVFV-SL-eGFP and iRVFV-ML, abundant expression of eGFP but also of Gn was observed within the same cells, suggesting that particles from both populations can co-infect individual cells and together provide a full set of genome segments encoding all the viral proteins. The co-expression of eGFP and Gn was accompanied by the detection of clusters of infected neighboring cells and a rapid increase in the percentage of infected cells over time, as was also observed after infection with the three-segmented RVFV-eGFP strain. This result strongly suggests that co-infection with complementing incomplete particles allows the rescue of a three-segmented virus. Notably, complementation by incomplete particles was efficient in both BSR-T7/5 and C6/36 cells even at low MOIs **(≥** 0.1 and **≥** 0.01, respectively) (**Fig. 4b-c**). Next, to confirm whether cells simultaneously exposed to the two different populations of incomplete particles produce infectious progeny able to spread efficiently, we passaged the supernatants of both BSR-T7/5 and C6/36 cells to naive cells. The results showed that early after infection (24 hpi), a high proportion of the cell population abundantly co-expressed eGFP and Gn, with clusters of positive cells rapidly expanding, suggesting that infectious virus was rescued (**Fig. 4d**).

**Fig. 4.**
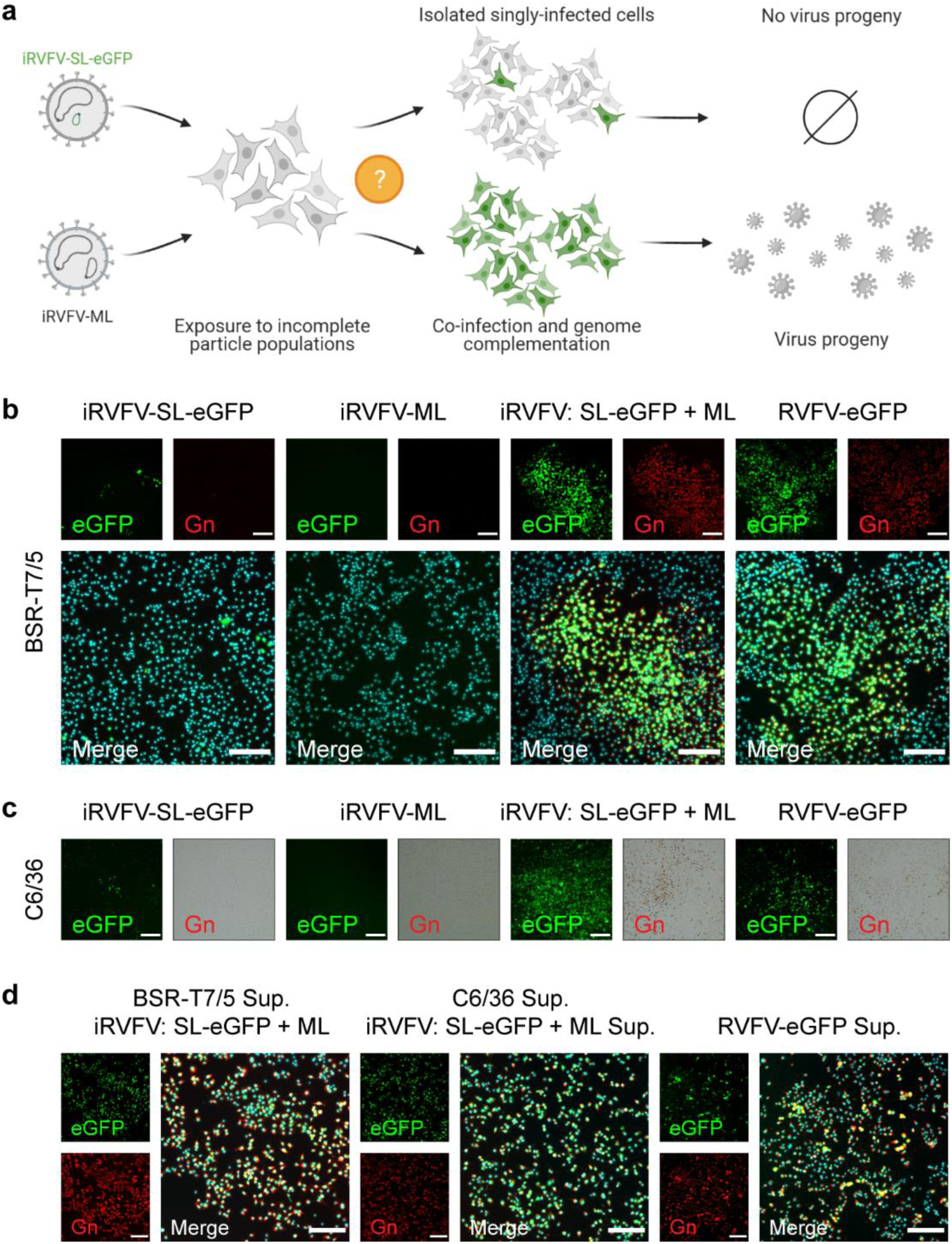
Co-infection with complementing incomplete RVFV particles results in efficient virus replication and spread among mammalian and insect cells. **a** Schematic representation of the individual or simultaneous exposure of mammalian (BSR-T7/5) and insect (C6/36) cells to non-spreading iRVFV-SL-eGFP and iRVFV-ML particles. If individual susceptible cells become co-infected by the two different incomplete particles, genome complementation could eventually allow virus replication and production of infectious progeny. **b, c** BSR-T7/5 cells (**b**) and C6/36 cells (**c**) were mock-infected, infected exclusively with one population of incomplete particles (MOI 0.1), or simultaneously infected with iRVFV-SL-eGFP and iRVFV-ML (MOI 0.1 for each virus). Cells infected with a three-segmented RVFV-eGFP (MOI 0.2) were used as a positive control. In all cases, infected cells were analyzed at 24 h (BSR-T7/5 cells) or 72 h (C6/36 cells) post-infection by following the expression of eGFP (green) via direct fluorescence microscopy and the expression of Gn (red) via an immunofluorescence assay in BSR-T7/5 cells or an immunoperoxidase monolayer assay in C6/36 cells. For detection of Gn in C6/36 cells, the immunoperoxidase monolayer assay was chosen because the fluorescent signal got drastically reduced upon fixation of the cells. Expression of Gn was detected with rabbit polyclonal anti-Gn serum in combination with Alexa Fluor 568-conjugated secondary antibodies (immunofluorescence assay) or with HRP-conjugated secondary antibodies (immunoperoxidase monolayer assay). Cell nuclei (cyan) were visualized with DAPI. Scale bars, 200 μm. Infections were also carried out at different MOIs (ranging from 0.001 to 0.25 for each virus, **Supplementary Fig. 1**). **d** Supernatants from BSR-T7/5 and C6/36 co-infected cells or cells infected with RVFV-eGFP were passaged onto naive BSR-T7/5 cells. Cells were examined at 24 h post-infection combining direct fluorescence microscopy and immunofluorescence as described for panels in **b**. Scale bars, 200 μm.

Besides tracking the course of infection at the cell population level, we next analyzed the intracellular genome content of individual infected BSR-T7/5 cells using a combined smFISH-immunofluorescence method^15^ (**Fig. 5a**). As expected, in cells exposed exclusively to iRVFV-SL-eGFP, we detected abundant copies of the S and L genome segments, while the M segment and the Gn glycoprotein were absent. No expression of eGFP or Gn was detected in cells exclusively exposed to iRVFV-ML particles, as the S genome segment is missing in this population, and thus no genome replication or transcription could take place. Contrary to cells exposed to only one population of incomplete particles, cells simultaneously exposed to both iRVFV-SL-eGFP and iRVFV-ML particles showed abundant copies of each of the three genome segments, similar to cells infected with the three-segmented RVFV-Clone 13 (**Fig. 5b**).

**Fig. 5.**
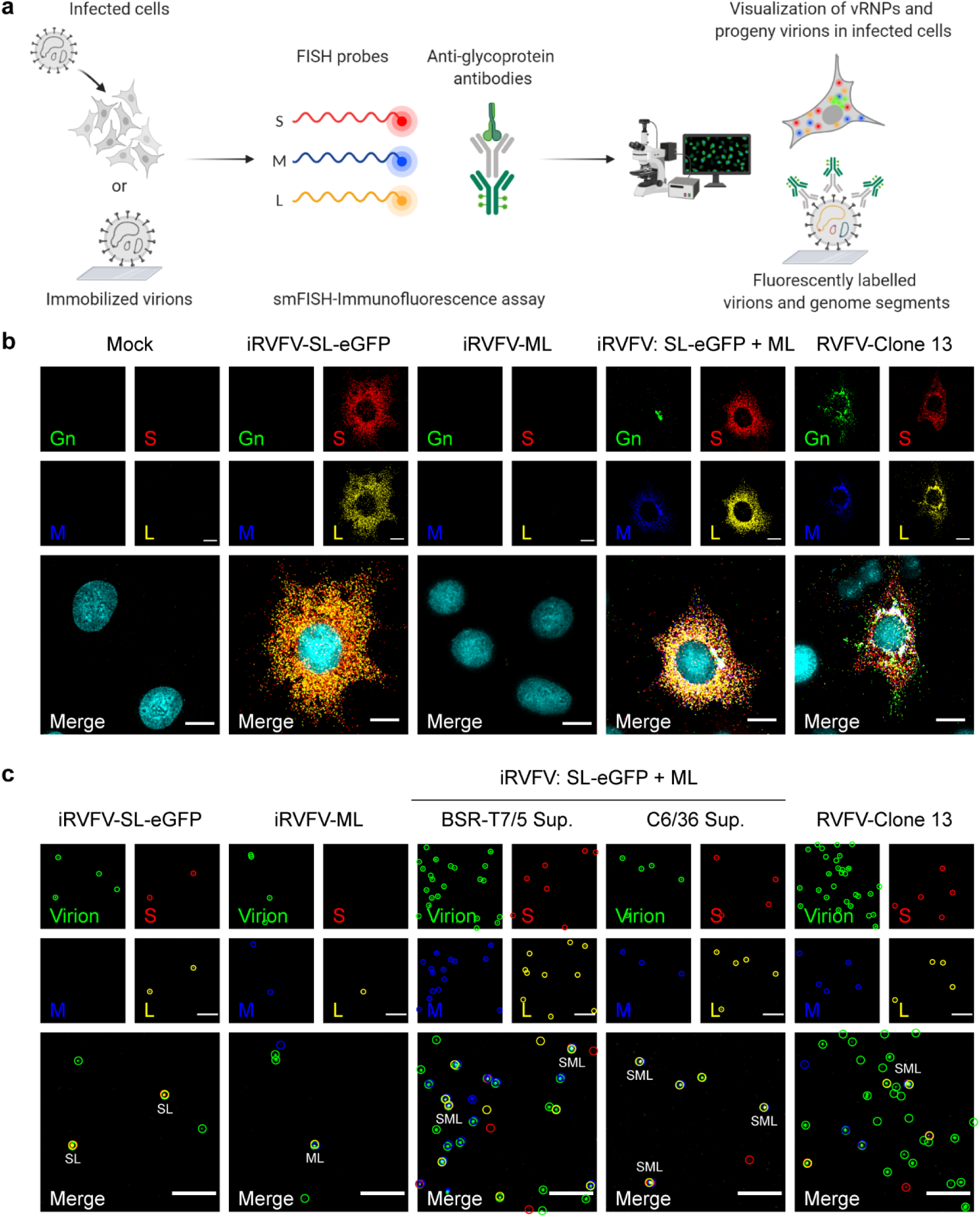
Visualization of viral genome segments in RVFV-infected cells and in immobilized RVFV virions from virus stocks and co-infection supernatants. **a** Schematic representation of the single-molecule viral RNA FISH-immunofluorescence method used to visualize the intracellular viral RNP composition of infected cells and the intra-virion genomic composition of virus stocks and culture supernatants (illustration based on ^15^). **b** BSR-T7/5 cells were mock-infected, infected exclusively with one population of incomplete particles (MOI 1), or simultaneously infected with iRVFV-SL-eGFP and iRVFV-ML (MOI 1 for each virus), and fixed at 16 h post-infection. Cells infected with a three-segmented RVFV-Clone 13 (MOI 1) were used as a positive control. **c** RVFV virions from incomplete particle stocks and supernatants of co-infected BSR-T7/5 and C6/36 cells were immobilized on coverglass by incubation for 5 h at 28°C. The S segment (N gene, red), M segment (polyprotein gene, blue) and L segment (RdRp gene, yellow) were hybridized using probe sets labeled with CAL Fluor Red 610, Quasar 670 and Quasar 570, respectively. RVFV particles (green) were detected with antibody 4-D4^38^ targeting the Gn glycoprotein in combination with Alexa Fluor 488-conjugated secondary antibodies. Cell nuclei (cyan) were visualized with DAPI. Merge images show the overlay of five (for cells) or four (for virions) individual channels. Individual spots, each representing either a single vRNP or a virus particle, were detected and assessed for co-localization in ImageJ with the plugin ComDet. Colored circles display the spots detected in each channel and their co-localization in the merge image. Scale bars, 10 μm (**b**), 5 μm (**c**).

Finally, we used our smFISH-immunofluorescence method to obtain additional insights into the intra-virion composition of the incomplete particle populations and the progeny virions released upon co-infection with these incomplete particles (**Fig. 5a**). Importantly, the M genome segment was not present in iRVFV-SL-eGFP particles, and the S genome segment was not present in iRVFV-ML particles, confirming the absence of the respective segment in each of the incomplete particle populations. Notably, just as in stocks of the three-segmented positive control RVFV-Clone 13, a fraction of the virions produced by co-infected cells was shown to contain the complete set of S, M and L genome segments, ultimately confirming the rescue of infectious three-segmented virus upon co-infection with incomplete particles (**Fig. 5c**).

### Co-infection with complementing incomplete RVFV particles supports *in vivo* virus spread in mosquitoes

Having confirmed the ability of incomplete RVFV particles to generate a productive infection *in vitro* upon co-infection, we hypothesized that incomplete bunyavirus particles could also play a role in between-host virus transmission. To investigate this, groups of *Aedes aegypti* mosquitoes were fed with bovine blood meals spiked with diverse virus populations. Mosquitoes were exposed to a mock blood meal (group #1), a blood meal spiked with a single incomplete virus particle population (group #2 to iRVFV-SL-eGFP and group #3 to iRVFV-ML), a blood meal spiked with three-segmented RVFV-mCherry2 (group #4) or a blood meal spiked with a mixture of incomplete particles (iRVFV-SL-eGFP and iRVFV-ML, group #5). After incubation for 12-15 days post-feeding at 28°C, mosquitoes were analyzed for virus replication in their bodies and dissemination to the saliva (**Fig. 6a**). To this end, body homogenates and saliva samples were tested by virus isolation on BSR-T7/5 cells, using the expression of virus-encoded eGFP or mCherry2 (depending on the virus) as a read-out. All the mosquitoes fed on a mock blood meal (group #1, n = 15) or a blood meal spiked with only one population of incomplete virus particles (group #2, n = 25; and group #3, n = 20) were virus-negative in their bodies and saliva (**Fig. 6b**). No virus spread was observed in these groups, as iRVFV-SL-eGFP and iRVFV-ML both lack one genome segment, either the M or the S segment. In group #4, 33 out of 52 (63%) mosquitoes fed on a blood meal spiked with three-segmented RVFV-mCherry2 were found virus-positive in their bodies, confirming the high vector competence of the mosquitoes for RVFV. Eleven of these 33 mosquitoes were also virus-positive in their saliva, implying that RVFV-mCherry2 had replicated in the mosquito midgut, disseminated throughout the body, reached the salivary glands and was excreted in saliva. Strikingly, 24 out of 62 (39%) mosquitoes in group #5, which were fed on a blood meal spiked with a mixture of iRVFV-SL-eGFP and iRVFV-ML particles, were found to have virus-infected bodies. Of those 24, three mosquitoes were found to be virus-positive in their saliva (**Fig. 6b-c**). These results show that genome complementation via co-infection with two populations of RVFV exclusively comprising incomplete particles can occur *in vivo* as well, supporting virus spread in mosquitoes and presumably contributing to virus transmission between hosts.

**Fig. 6.**
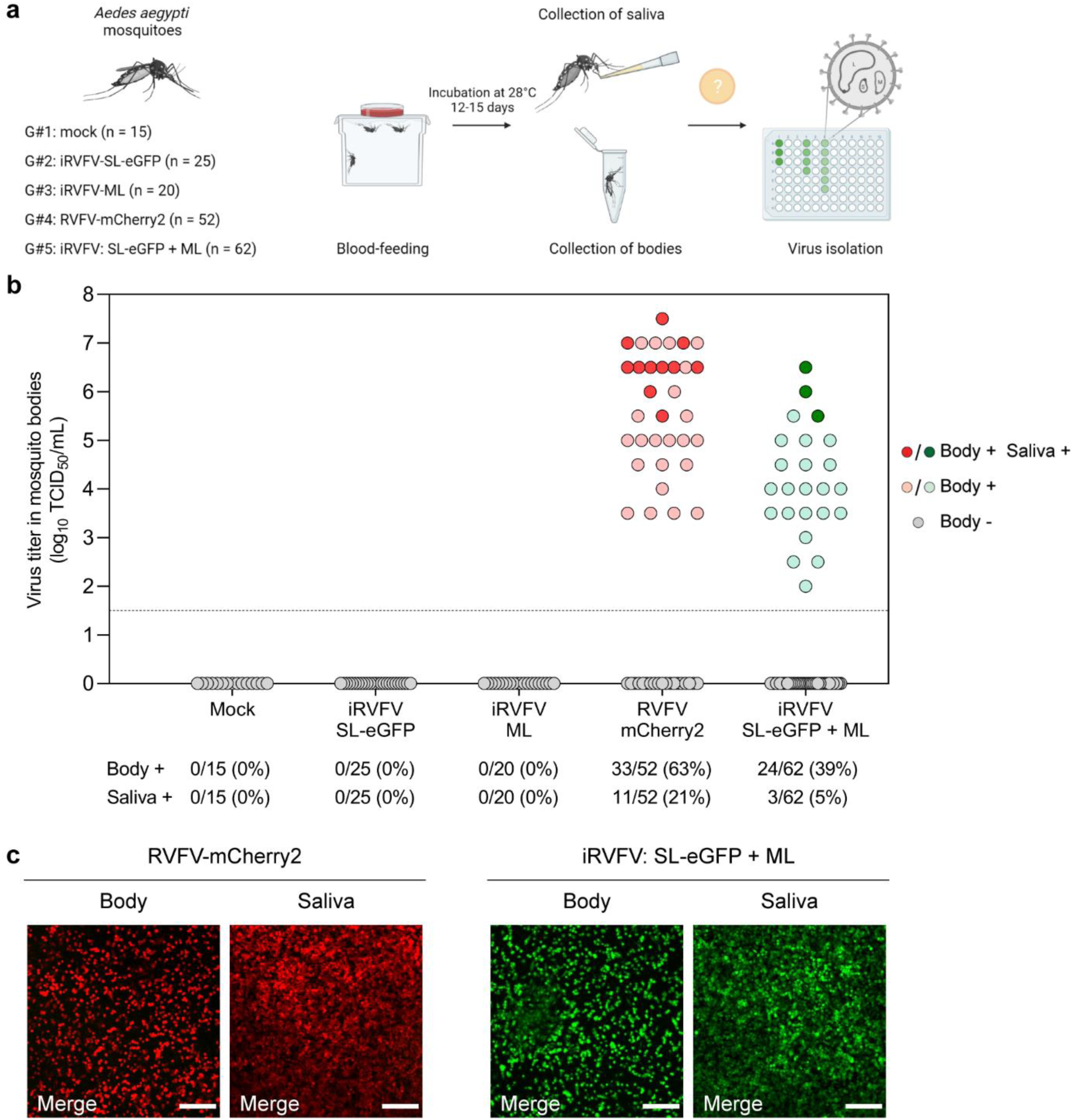
RVFV incomplete particles complement upon co-infection and allow virus replication and spread in mosquitoes. **a** Schematic representation of the experimental design. Five groups of *Aedes aegypti* mosquitoes (n = 15-62 per group) were fed a blood meal spiked with different RVFV preparations and housed at 28°C. After 12-15 days, mosquitoes were sedated with CO_2_ and body and saliva samples were collected. RVFV infection of mosquito bodies and transmission to mosquito saliva were assessed via virus isolation with a fluorescence microscopy read-out. **b** RVFV infectious titers in mosquito bodies. Dots represent individual mosquitoes and are color-coded gray for negative bodies, light red (group #4) or light green (group #5) for positive bodies, and solid red (group #4) or solid green (group #5) for both body and saliva virus-positive mosquitoes. The dashed line indicates the limit of detection (10^1.5^ TCID_50_/mL). The incidence (in absolute numbers and percentages) of RVFV infection (bodies) and transmission (saliva) is indicated on the bottom for each group. Data corresponding to mosquitoes from groups #4 and #5 derive from two independent experiments. **c** Representative fluorescence microscopy images of BSR-T7/5 cells inoculated with virus-positive body and virus-positive saliva samples from groups #4 (RVFV-mCherry2) and #5 (iRVFV: SL-eGFP + ML) at 48 h post-infection. Merge images show the overlay of two individual channels (eGFP and mCherry2). Scale bars, 200 μm.

### Predicting the contribution of incomplete particles to virus spread

We showed that experimental co-infection with complementing two-segmented incomplete particles results in rescue of spreading three-segmented virus both *in vitro* and *in vivo*. However, incomplete particles naturally co-exist within a mixed population of empty, incomplete and complete particles. To study to what extent incomplete particles actually contribute to virus spread, we modeled and compared RVFV infection dynamics in three different scenarios: (i) non-selective genome packaging without productive co-infection by incomplete virus particles, (ii) non-selective genome packaging with productive co-infection by incomplete virus particles and (iii) selective genome packaging (**Fig. 7a**). In both the first and second scenarios, there is non-selective packaging, and the distribution of genome segments over virus particles approximately follows the empirical distribution. In the first scenario, the model assumes that the infection is driven exclusively by complete three-segmented particles, which only account for a small fraction of the particle population. By contrast, the second scenario allows the infection to progress by both complete particles and co-infecting incomplete particles, provided all segments are present in a single cell. Lastly, in the third scenario, it is assumed that all the particles are complete due to selective packaging, which serves as a reference point. Since there are differences in genome packaging efficiencies between mammalian and insect hosts^15^, and therefore the exact ratio between empty, incomplete and complete particles differs considerably depending on the host, we performed simulations with RVFV populations representative for those found in mammalian and insect hosts. Model parameters were defined to represent the conditions of localized virus spread (*i*.*e*. mean-field simulations in a small number of cells, *e*.*g*. 100), and the model predicts how the number of infected cells changes over time. In the first scenario, the number of infections increases the slowest, being limited by the small number of complete virus particles. In the second scenario, when the model takes into account that incomplete particles can generate a productive infection through complementation, the number of infected cells increases more rapidly, highlighting the contribution of the incomplete particles to virus spread. However, the number of infected cells does not increase as quickly as in the third scenario in which all virus particles are complete, suggesting there is still a cost to non-selective packaging. Importantly, the model is sensitive to the host in which the virus is replicating, suggesting that the contribution of incomplete particles to virus spread is greater in mammalian cells compared to insect cells, due to the smaller fraction of complete particles in mammalian-derived virus.

**Fig. 7.**
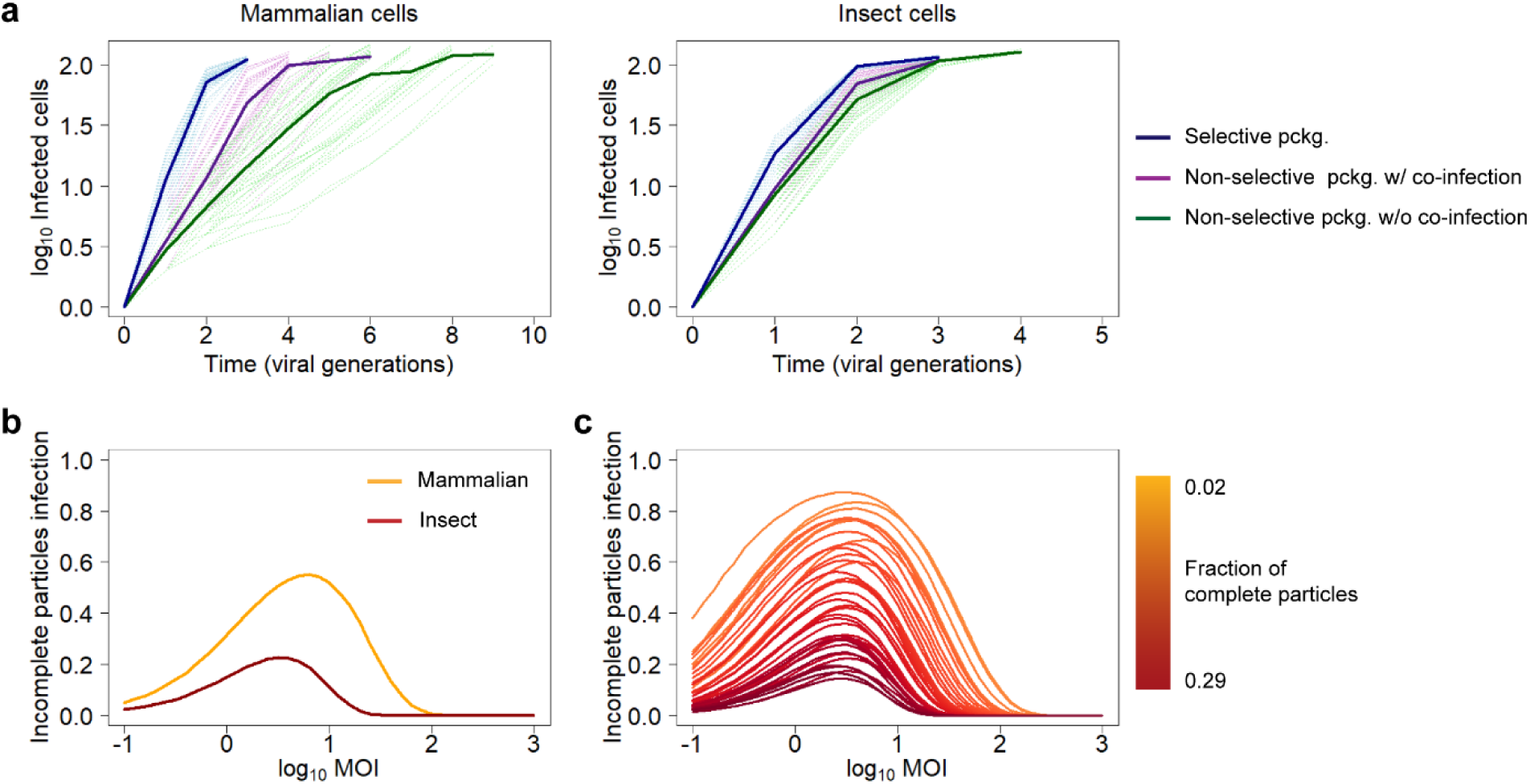
Predicting the contribution of incomplete particles to within-host virus spread. **a** Predicted RVFV infection dynamics in mammalian and insect cells in three different scenarios: selective genome packaging, non-selective genome packaging without co-infection by incomplete particles, and non-selective genome packaging with co-infection and productive complementation by incomplete particles. The low number of cells in the model is representative of localized virus spread. Solid lines represent the mean of n = 50 simulations per scenario and the light-colored dashed lines represent individual simulations. pckg.: packaging. **b** Contribution of incomplete particles to virus spread in mammalian and insect cells as a function of the MOI in a model representative of non-selective genome packaging allowing co-infection by incomplete particles. Lines represent the mean of n = 1 × 10^5^ – 2 × 10^6^ simulated cells, with a larger number of cells for the low MOI conditions where overall infection frequencies are low. **c** Contribution of incomplete particles to virus spread as a function of the MOI for starting inoculums with different frequencies of complete particles. Lines are color-coded in a gradient of dark yellow to dark red based on an increasing fraction of complete particles (ranging from 0.02 to 0.29) within the virus population.

To account for the contribution of incomplete particles in a model of localized virus spread, we next investigated the relationship between this contribution and the MOI. To this end, we calculated the fraction of infected cells comprising a full set of genome segments as a result of co-infection with incomplete particles over a range of MOIs (**Fig. 7b**). The simulations show that the MOI has a pronounced effect on the contribution of incomplete particles to virus spread, with a limited contribution at extreme MOIs. At low MOIs there is a low probability of particles co-infecting the same cell and consequently complementation events are rare, whereas at high MOIs in most cells infection will be initiated by at least one complete particle and complementation of incomplete particles is not required for virus spread. Notably, at intermediate MOIs, infection initiated by complementing incomplete particles is more common and plays an important role in virus spread. Here again, the contribution of incomplete particles is estimated to be higher in the mammalian host compared to the insect host, because insect-derived virus preparations contain a larger fraction of complete particles^15^. To gauge how general these model results are, we considered the relationship between MOI and the contribution of incomplete virus particles over a broad range of randomly selected genome segments frequencies for a three-segmented virus. Interestingly, the larger contribution of incomplete particles at intermediate MOIs appears to be a general feature over different virus particle compositions (**Fig. 7c**). In line with what we observed for the mammalian- and insect-derived virus, this contribution becomes less relevant when the fraction of complete particles becomes larger.

## Discussion

Mammalian-infecting viruses with an incomplete genome are unable to spread autonomously and have principally been considered as a source of interference in the course of an infection^17^. Segmented viruses with a non-selective genome packaging strategy are prone to generate large fractions of empty and incomplete particles^14,15^, which in theory could detriment fitness of the virus population as a whole. We hypothesized, however, that incomplete bunyavirus particles may contribute to virus spread by complementing their genomes inside cells co-infected with multiple particles, resembling the strategy that multipartite viruses use to replicate in plants and fungi. Using the three-segmented RVFV as a prototype bunyavirus with a non-selective genome packaging strategy, we show here that incomplete bunyavirus particles can contribute significantly to within-host spread and between-host transmission.

By creating fluorescently labeled RVFV variants, we were able to visualize infection of single cells by more than one virus particle and to show that mammalian and insect cells are prone to co-infection as a function of the MOI. Notably, the fractions of experimentally infected and co-infected cells coincided with theoretical models that predict the absence of a major bottleneck determining the probability of (co-)infection other than intrinsic heterogeneous susceptibility within a host cell population. In our experiments, we exposed cells to two different RVFV variants simultaneously, as this represents a common scenario during a localized infection, in which non-infected cells are exposed to the burst of virus particles released by a neighboring infected cell. Nevertheless, it should be noted that susceptibility to co-infection may differ when cells are exposed to the viruses at different time points due to innate immune responses and/or super-infection exclusion mechanisms, as has already been shown for other arthropod-borne viruses^18–21^. The frequency of co-infection events upon delayed exposure to a second virus is thus likely lower compared to simultaneous exposure. Related to this, it would be interesting to compare host cell responses following infection by a three-segmented virus, a two-segmented virus capable of replicating its genome, and a two-segmented virus unable to replicate its genome.

By generating two different incomplete virus particle populations entirely dependent on co-infection (iRVFV-SL-eGFP and iRVFV-ML), we showed at the cell population level and at single-cell level that within-host genome complementation occurs readily in mammalian and insect cells, resulting in infectious progeny able to spread in a similar way as progeny resulting from infection with a virion containing all three genome segments. Importantly, by examining the genome composition of individual virions, we confirmed that cells simultaneously exposed to complementing incomplete particle populations can produce complete virus particles containing all three genome segments. As expected, the contribution of genome complementation to virus spread was found to depend on the MOI. Specifically, at both low and very high MOIs, the contribution to virus spread by incomplete particles was predicted to be negligible. The former is explained by the low probability of co-infection and the latter by the high probability of infection by a complete particle. Experimentally, we confirmed that at very low MOIs (≤ 0.01 for BSR-T7/5 cells and ≤ 0.001 for C6/36 cells), the chance of a co-infection event is rather low and consequently the probability of a productive infection is reduced (**Supplementary Fig. 1**). These results are in line with the mathematical models that predicted the contribution of incomplete particles to be highest at intermediate MOIs.

A previous report on the eight-segmented influenza A virus showed that a virus entirely dependent on co-infection replicated efficiently in guinea pigs but was less infectious than the wild-type virus and was not able to transmit between animals^22^. Here, we show that *in vivo* bunyavirus genome complementation can occur in the mosquito vector. It should be noted that after a mosquito ingests an infectious blood meal, the resulting MOI is probably very low, which reduces the probability of multiple particles entering the same cell. Despite the presumably low MOI, a fraction of the mosquitoes given a blood meal spiked with virus particles exclusively containing incomplete sets of genomes were found to be infected (both in bodies and saliva). Infected mosquito bodies can only be explained by complementing particles co-infecting the same midgut cell followed by the generation of a mixed virus progeny including three-segmented virus. Most likely, these co-infections occur in a small number of midgut epithelial cells that are highly susceptible to the virus, as reported previously for other arthropod-borne viruses^23–26^. Furthermore, the rescue of infectious virus from mosquito saliva indicates that the virus progeny was able to disseminate from the midgut cells to the hemocoel, generally considered a major bottleneck for virus dissemination in mosquitoes ^27^, and from the hemocoel to the salivary glands. Due to the absence of complete particles in the starting inoculum of the iRVFV-SL-eGFP and iRVFV-ML mixture, it was expected that the infectivity of this mixture was lower than the infectivity of the three-segmented RVFV-mCherry2. Although with lower efficiency, the fact that we rescued infectious virus from the saliva of mosquitoes fed with the mixture of incomplete particles suggests that incomplete particles play a role in between-host virus transmission. Interestingly, we observed a positive correlation between the infectious virus titer in the body and the presence of infectious virus in saliva, with virus-positive saliva samples corresponding to mosquitoes with a high virus titer in their bodies.

Several mechanisms have been proposed that may compensate for the fitness cost resulting from inefficient bunyavirus genome packaging. For instance, the incorporation of more than three genome segments per particle, which increases the likelihood of incorporating a complete set of segments^13^. Alternatively, groups of virions could be transmitted together in structures known as collective infectious units, jointly delivering multiple virions to target cells and drastically increasing the local MOI^28–31^. The latter, together with our finding that incomplete particles can genetically complement within co-infected cells, has important biological implications particularly for within-host virus spread, where following a primary infection of a target organ, neighboring cells become a local environment exposed to intermediate and high MOIs. In this study, we focused on co-infections by incomplete particles of the same virus. However, co-infections by incomplete particles of different but genetically related viruses could give rise to reassortment events, adding to the potential roles of incomplete particles in the infection cycle and evolution of bunyaviruses.

Altogether, the results of this study show that upon co-infection, incomplete bunyavirus particles can initially drive efficient progeny virus generation and spread, even in the absence of three-segmented RVFV infectious particles in the inoculum. In the context of natural infection with a mixed bunyavirus particle population, we propose that incomplete particles, instead of interfering, can substantially contribute to within-host spread and between-host transmission, facilitating the dual life cycle of bunyaviruses in mammalian and insect hosts (**Fig. 8a-b**). Moreover, we propose that this contribution, particularly important at intermediate MOIs (**Fig. 8c**), partially compensates for the disadvantage of inefficient genome packaging. In conclusion, the present work stands out an important role of incomplete particles in the infection cycle of bunyaviruses, revealing previously unrecognized parallels in the life cycles of segmented viruses and multipartite viruses.

**Fig. 8.**
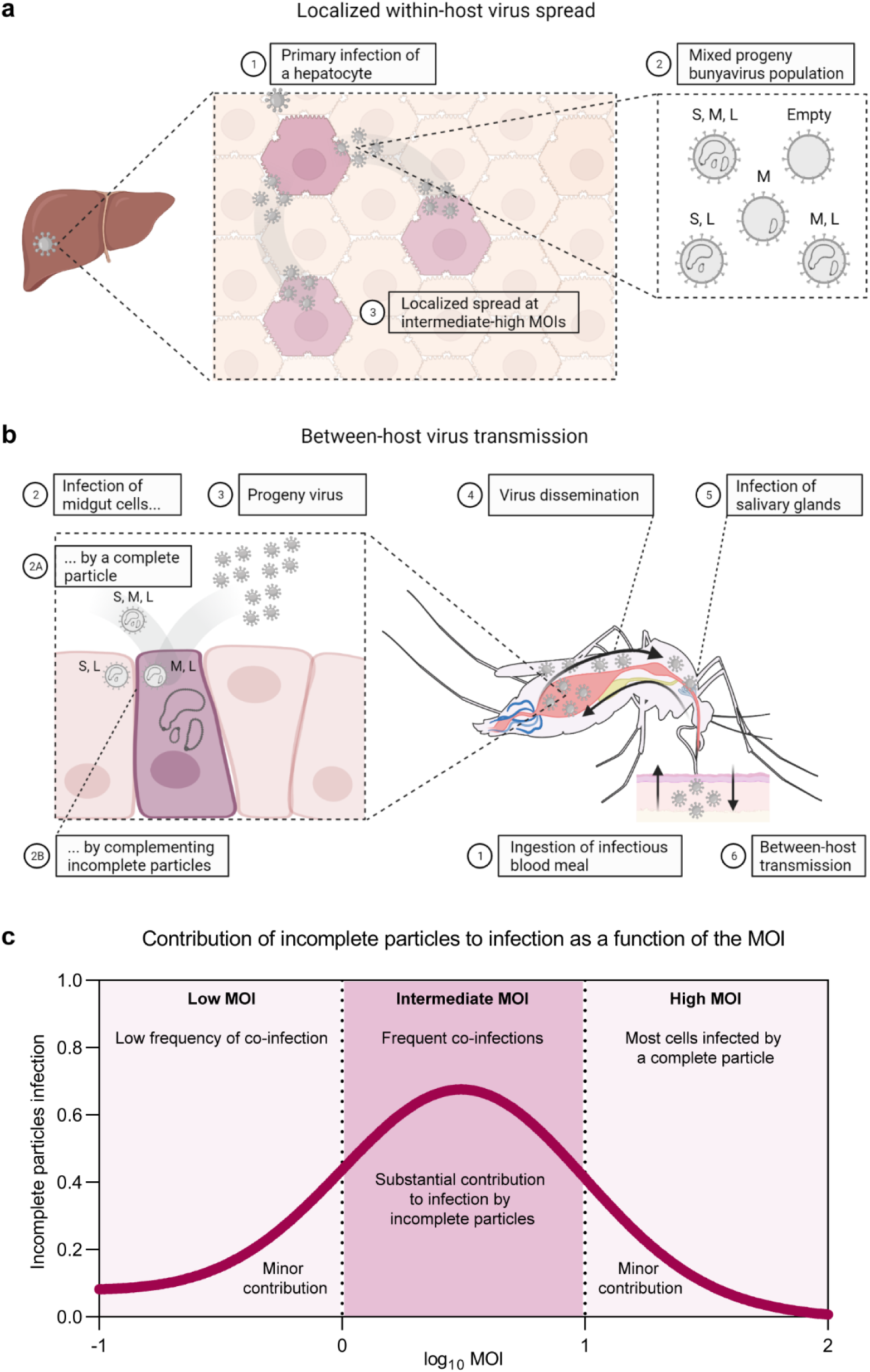
Schematic representation of the proposed model describing the contribution of incomplete virus particles to within-host spread and between-host transmission. **a** Localized within-host virus spread. (1) Most likely, a complete three-segmented particle derived from an infected mosquito vector starts a primary infection in a target organ (*e*.*g*. liver). (2) Due to a non-selective genome packaging strategy, bunyavirus-infected cells give rise to a virus progeny population comprising a mixture of complete, incomplete and empty virus particles. Although non-infectious on their own, incomplete particles with a complementing genome can co-infect the same cell and generate infectious progeny, contributing to localized secondary virus spread. (3) At the organ and tissue levels, the intermediate-high local MOI can greatly facilitate the occurrence of this phenomenon. **b** Between-host virus transmission. (1) The life cycle of bunyaviruses involves replication in both mammalian and insect hosts. For transmission, mosquitoes ingest an infectious blood meal from an infected animal. (2) Either through a complete three-segmented particle (2A) or through co-infection by complementing incomplete particles (2B), the virus infects a small group of highly susceptible cells of the mosquito midgut and (3) generates infectious progeny. (4) The virus disseminates from the midgut into the hemocoel and (5) reaches the salivary glands, where the virus also replicates. (6) To complete the cycle, the virus is excreted in the mosquito saliva and can be transmitted to a mammalian host through a mosquito bite (illustration inspired on ^27^). Although perhaps at a lower frequency than in a localized environment, co-infection with incomplete particles can contribute to between-host virus transmission. Viruses transmitted as collective infectious units can facilitate the co-infection of a single cell by multiple particles. **c** Conceptual illustration of the relationship between the MOI and infections caused entirely by incomplete virus particles. At low MOIs (< 1), infections caused by co-infection with incomplete particles are rare because on average less than a single virus particle will enter each cell. This situation is likely to represent the conditions of between-host transmission and early infection stages in a new host, where our data show that although incomplete particles have reduced infection potential, infection is still possible. At intermediate MOIs (1-10), infection via co-infecting incomplete particles becomes more common, as on average more virus particles enter each cell. This situation is likely to be representative of later infection stages within a host, when cells are exposed to larger numbers of virus particles. At high MOIs (> 10), infection caused only by incomplete virus particles becomes rare as most cells will also be infected by a complete virus particle. In this regime, incomplete particles can still contribute genetically to infection, but complementation between virus particles is no longer needed to establish infection. This latter situation may not be representative of conditions found in natural infections but can be generated in cell culture.

## Methods

### Cell lines

BSR-T7/5 cells (Golden hamster kidney, CVCL_RW96) stably expressing bacteriophage T7 RNA polymerase^32^ were maintained in Glasgow minimum essential medium (G-MEM) supplemented with 5% fetal bovine serum (FBS), 1% antibiotic/antimycotic, 1% MEM non-essential amino acids (MEM NEAA) and 4% tryptose phosphate broth at 37°C and 5% CO_2_. For stable maintenance of the cell line, the medium was supplemented with 1 mg/mL Geneticin (G-418 sulfate) every other passage. Vero E6 cells (African green monkey kidney, ATCC CRL-1586) were maintained in MEM supplemented with 5% FBS, 1% antibiotic/antimycotic, 1% MEM NEAA and 2 mM L-glutamine at 37°C and 5% CO_2_. C6/36 cells (*Aedes albopictus* larva, ATCC CRL-1660) were maintained in L-15 medium (Leibovitz) (Sigma-Aldrich) supplemented with 10% FBS, 1% antibiotic/antimycotic, 1% MEM NEAA and 2% tryptose phosphate broth at 28°C. Cell culture media and supplements were purchased from Gibco, unless specified otherwise.

### Viruses

Virus stocks of RVFV strain Clone 13^33^ were obtained following infection of Vero E6 cells at an MOI of 0.005. Recombinant fluorescently marked variants of RVFV strain 35/74 (accession numbers JF784386-88)^34^, expressing either eGFP (RVFV-eGFP) or mCherry2 (RVFV-mCherry2), were generated by a pUC57 transcription plasmid-based reverse genetics system^16^. Briefly, BSR-T7/5 cells were seeded in 6-well cell culture plates at 1.5-2.0 × 10^5^ cells/well (2 mL volume) and incubated for 18-24 h to generate a sub-confluent monolayer. Cells were subsequently co-transfected with the plasmids pUC57_S-FP (where FP corresponds to either eGFP or mCherry2 in place of the NSs gene), pUC57_M and pUC57_L, encoding the RVFV-35/74 S, M and L genome segments, respectively, in antigenomic-sense orientation. Transfections were performed using *Trans*IT-LT1 transfection reagent (Mirus), with slight modifications to the manufacturer’s instructions: 3-5 h prior to transfection, the seeding supplemented G-MEM was removed and substituted by 1 mL of Opti-MEM I Reduced-Serum medium. For transfection, 2500 ng total plasmid/well (equal amounts per plasmid) with a 1:4 plasmid to transfection reagent ratio were used. At 3-5 h post-transfection, 1 mL of supplemented G-MEM was added. At 3 days post-transfection, culture supernatants were harvested and clarified by low-speed centrifugation. High-titer virus stocks of RVFV-eGFP and RVFV-mCherry2 were obtained following infection of BSR-T7/5 cells at an MOI of 0.001.

### Generation of incomplete virus particles

Incomplete RVFV particles lacking either the S segment (iRVFV-ML) or the M segment (iRVFV-SL-FP) were generated by reverse genetics. For the production of iRVFV-ML, BSR-T7/5 cells were co-transfected with pUC57_M, pUC57_L and the protein expression plasmid pCAGGS_N (encoding for a codon-optimized RVFV-35/74 nucleocapsid protein). For the production of iRVFV-SL-FP, BSR-T7/5 cells were co-transfected with pUC57_S-FP (where FP corresponds to eGFP or mCherry2 in place of the NSs gene), pUC57_L and the protein expression plasmid pCAGGS_NSmGnGc (encoding for the RVFV-35/74 polyprotein). Transfections were performed using *Trans*IT-LT1 transfection reagent (Mirus), following the protocol described for the generation of recombinant fluorescently marked RVFV-35/74 variants.

To obtain a high-titer stock of iRVFV-ML particles, the rescued supernatant from the transfection was concentrated by ultracentrifugation at 28000 rpm, 4°C for 2 h (SW 32 Ti swinging-bucket rotor, Optima LE-80K Beckman Coulter) using a 25% w/v sucrose cushion. Furthermore, to obtain higher titers of iRVFV-SL-FP particles, cells initially transfected with the three plasmids were repeatedly transfected two additional times with the protein expression plasmid pCAGGS_NSmGnGc, resulting in a polyclonal cell line replicating the S and L viral genome segments. New stocks of iRVFV-SL-FP particles were generated by a final transfection of the polyclonal cell line with pCAGGS_NSmGnGc. The latter transfections were performed using jetPEI transfection reagent (Polyplus) with a 1:3 plasmid to transfection reagent ratio. At 1 day post-transfection, culture supernatants were harvested and clarified by low-speed centrifugation.

### Virus titration

Infectious titers of rescued virus and virus stocks were determined in an end-point dilution assay in combination with either an immunoperoxidase monolayer assay (IPMA, as described below) or direct microscopy detection of fluorescent protein expression. BSR-T7/5 (3 × 10^4^ cells/well) or C6/36 (6 × 10^4^ cells/well) monolayers were incubated with 10-fold serial dilutions (starting at 1:10) of the samples for 72 h at 37°C and 5% CO_2_ (BSR-T7/5) or 28°C (C6/36). Samples were analyzed in triplicate or quadruplicate, and the titer was calculated as the median tissue culture infectious dose (TCID_50_/mL) using the Spearman-Kärber method.

### Growth curves of fluorescent virus variants

BSR-T7/5 monolayers (3 × 10^6^ cells/T75 flask) were infected with RVFV-eGFP or RVFV-mCherry2 at an MOI of 0.01. At 2 h post-infection, the inoculum was removed, cells were washed with PBS, and fresh medium was added. Cell culture supernatants were harvested at 0, 2, 12, 24, 48 and 72 h post-infection and were clarified by centrifugation at 2500 rcf for 10 min. Virus titers of the clarified supernatants were determined with an end-point dilution assay (fluorescence microscopy read-out) as described above. Growth curve determinations were performed with three biological replicates per time point.

### Genome segment-specific quantitative RT-PCR

As a conventional end-point dilution assay is not suited for determining the titer of non-replicating particles, we estimated the titer of iRVFV-ML stocks through genome segment-specific quantification via RT-qPCR, followed by a comparison with the genome copies of iRVFV-SL-eGFP and three-segmented RVFV-eGFP preparations with known infectious titer determined with the end-point dilution assay. Total RNA extractions of 80 μL of virus stocks lyzed with 240 μL of TRIzol LS Reagent (Invitrogen) were performed in triplicate with the Direct-zol RNA MiniPrep kit (Zymo Research), largely according to the manufacturer’s instructions, except for a more thorough in-column DNase I treatment. Namely, lyzed preparations were treated with 60 units of DNase I for 30 min. The extended DNase I treatment ensured the complete removal of residual plasmid DNA from the transfection. Total RNA was eventually eluted in 25 μL of DNase/RNase-free water (Zymo Research). Subsequently, viral cDNA was synthesized with the SuperScript IV First-Strand Synthesis System for RT-PCR (Invitrogen) using 100 units of SuperScript IV reverse transcriptase and a combination of S, M and L segment-specific primers (**Supplementary Table 1**). After the reverse transcription reaction, quantitative PCR amplifications were performed with the Power SYBR Green PCR Master Mix using 5 μL of 50-, 250- or 500-fold diluted cDNA preparations in a total volume of 25 μL, in combination with a 7500 Fast Real-Time PCR System (Applied Biosystems). Fragments from the three viral genome segments were amplified using specific primers (**Supplementary Table 2**) under the following conditions: an initial denaturation step at 95°C for 10 min; 40 cycles of denaturation at 95°C for 15 s, annealing at 59°C for 30 s and extension at 72°C for 36 s; and a single cycle of denaturation at 95°C for 15 s, annealing at 60°C for 1 min, denaturation at 95°C for 15 s and annealing at 60°C for 15 s. Per sample, an additional reaction intended to detect residual plasmid DNA was carried out using primers designed to amplify a fragment of the ampicillin resistance gene (*ampR*) (**Supplementary Table 2**) present in pUC57 and pCAGGS plasmids used for generating the different RVFV variants. Data were acquired and analyzed with the 7500 Fast System software version 1.4. (Applied Biosystems). Genome copies of each viral segment were calculated by intrapolation of the respective standard curve prepared with 10-fold serial dilutions of the viral segment cloned in pUC57 plasmids starting at 0.1 ng/μL.

**Table 1.** Description and mean titers of the virus stocks used for blood meals in the mosquito experiment.

| Group | Virus stock | Blood-virus mixture titer ($\log_{10}$ TCID<sub>50</sub>/mL) |
| --- | --- | --- |
| Group #1 | Mock | --- |
| Group #2 | iRVFV-SL-eGFP | 8.1 |
| Group #3 | iRVFV-ML | Not determined* |
| Group #4 | RVFV-mCherry2 | 8.1 |
| Group #5 | iRVFV-SL-eGFP + iRVFV-ML | 7.0 |

\*Virus stock prior to mixing with blood had an initial titer of $10^{7.1}$ TCID<sub>50</sub>/mL

### Flow cytometry

BSR-T7/5 cells were simultaneously co-infected with iRVFV-SL-eGFP and iRVFV-SL-mCherry2 at increasing MOIs (ranging from 0.1 to 2.5 for each virus). At 48 h post-infection, cells were trypsinized, spun down at 200 rcf for 5 min, fixed with 4% paraformaldehyde for 15 min, washed twice with PBS and resuspended in FACS sample buffer (PBS supplemented with 1% bovine serum albumin and 0.01% sodium azide). Mock-infected cells and cells infected exclusively with iRVFV-SL-eGFP or iRVFV-SL-mCherry2 at an MOI of 0.5 were used as negative and singly-infected controls, respectively. Flow cytometry was performed using a BD FACS Aria III (BD Biosciences) equipped with a standard laser and filter configuration. Cell subpopulations were categorized and quantified based on the expression of eGFP, mCherry2, both or none. Per sample, 50,000 events were recorded. Data was analyzed with FlowJo v10.7.1. The gating strategy applied involved discriminating cells from debris, followed by selection of single cells and assessment of fluorescent protein expression (**Supplementary Fig. 2**).

### Co-infections with incomplete virus particles

BSR-T7/5 (5 × 10^4^ cells/well) or C6/36 (1 × 10^5^ cells/well) monolayers seeded in 24-well cell culture plates were simultaneously co-infected with iRVFV-SL-eGFP and iRVFV-ML particles at increasing MOIs (ranging from 0.001 to 0.25 for each virus). At 2 h post-infection, the inoculum was removed, cells were washed with PBS, and fresh medium was added. At defined time points post-infection (varied per cell line and ranged from 24 h to 72 h), cells were examined for expression of eGFP via direct fluorescence microscopy. Expression of Gn in BSR-T7/5 and C6/36 infected cells was examined via an IPMA or an immunofluorescence assay (as described below). The expression of eGFP was followed over the course of the infection as an indicator of viral spread. Co-infections at the different MOIs were performed with three biological replicates. Cells infected exclusively with either iRVFV-SL-eGFP or iRVFV-ML particles were used as singly-infected controls, whereas cells infected with a three-segmented RVFV-eGFP (MOIs ranging from 0.002 to 0.5) were used as positive control for virus spread. Mock-infected samples were used as negative control.

### Immunostainings

BSR-T7/5 and C6/36 cells infected with different RVFV variants were subjected to an IPMA or an immunofluorescence assay to detect the expression of the viral proteins Gn and N. At defined time points post-infection (varied per experiment), cells were fixed with 4% paraformaldehyde for 15 min, washed with PBS supplemented with 0.5% Tween 80 (PBST), and permeabilized with 1% Triton X-100 in PBS for 5 min. Next, samples were blocked with 5% horse serum in PBS and subsequently incubated in sequential steps with primary and secondary antibodies diluted in blocking solution (**Supplementary Table 3**). Gn was detected with a polyclonal serum from a Gn-immunized rabbit (1:500 dilution, Thermo Fisher) as primary antibody. For the IPMA, HRP-conjugated goat polyclonal anti-rabbit immunoglobulin (1:500 dilution, P0448 Dako) was used as secondary antibody. For the immunofluorescence assay, goat polyclonal anti-rabbit IgG labeled with FITC (1:250 dilution, sc-2012 Santa Cruz Biotechnology) or donkey polyclonal anti-rabbit IgG labeled with Alexa Fluor 568 (1:500 dilution, A10042 Invitrogen) were used as secondary antibodies. N was detected with a monoclonal mouse hybridoma (1:100 dilution, F1D11 CISA-INIA) as primary antibody and a goat polyclonal anti-mouse IgG labeled with Alexa Fluor Plus 488 (1:500 dilution, A32723 Invitrogen) as secondary antibody. Incubations with the blocking solution, primary and secondary antibodies were each for 1 h at 37°C. Plates were washed with PBST after permeabilization, between the addition of primary and secondary antibodies, and prior to staining. In the IPMA, a 0.2 mg/mL amino ethyl carbazole solution in 500 mM acetate buffer pH 5.0, 88 mM H_2_O_2_ was added as substrate. In the immunofluorescence assay, cell nuclei were stained by incubation with 1 μg/mL DAPI in PBS for 5 min. Mock-infected samples and samples without the addition of primary antibodies were used as negative controls.

### Single-molecule RNA FISH-immunofluorescence

Experiments were performed with slight modifications to the Stellaris protocol for simultaneous FISH-immunofluorescence in adherent cells (Biosearch Technologies)^35–37^. BSR-T7/5 monolayers (1.5 × 10^4^ cells/well) were seeded on CultureWell 16 removable chambered coverglass (Grace Bio-Labs) and allowed to attach for at least 2 h at 37°C and 5% CO_2_. Cells were infected with the different RVFV variants at an MOI of 1. At 2 h post-infection, the inoculum was removed and the medium was refreshed. At 16 h post-infection, cells were fixed and permeabilized with a 3:1 mixture of methanol (Klinipath)-glacial acetic acid (Merck) for 10 min. Cells were subsequently washed twice with PBS and once with prehybridization buffer (10% deionized formamide [Millipore] in 2x concentrated SSC [Gibco]) for 5 min. Cells were then incubated for 4-16 h at 37°C with 100 μL/well of virus-specific FISH probe sets (**Supplementary Table 4**) and primary antibodies in hybridization buffer (10% deionized formamide, 10% dextran sulfate [Sigma-Aldrich], 2 mM vanadyl ribonucleoside complexes [VRC, Sigma-Aldrich] in 2x SSC). FISH probes were added at a final concentration of 250 nM. Gn was detected with hybridoma 4-D4^38^ supernatant (1:160 dilution) as primary antibody (**Supplementary Table 3**). Following hybridization and incubation with primary antibodies, cells were extensively washed at 37°C (twice with pre-hybridization buffer for 30 min and twice with 2x SSC for 15 min). Subsequently, cells were incubated with 100 μL/well of secondary antibody for 1 h at 37°C. A goat polyclonal anti-mouse IgG labeled with Alexa Fluor 488 (1:1000 dilution, A-11001 Invitrogen) was used as secondary antibody (**Supplementary Table 3**). Next, cells were washed twice with 2x SSC, and nuclei were stained by incubation with 100 μL/well of 1 μg/mL DAPI in 2x SSC for 5 min. Finally, cells were washed with 2x SSC and submerged in VectaShield antifade mounting medium H-1000 (Vector Laboratories). For analysis of virus stocks, undiluted or 1:3 diluted virus stocks were added on CultureWell 16 removable chambered coverglass and virions were allowed to attach to the surface for 5 h at 28°C. The same procedure as described for adherent cells was followed from the fixation step onwards. The specificity of the FISH probes and antibodies was reported previously^15^. Mock-infected samples and samples without primary antibodies were used as negative controls.

### Image acquisition and analysis

Light microscopy images were acquired with a Leica Model DMi1 inverted microscope and 10x HI PLAN I or 20x HI PLAN I objectives. Fluorescence microscopy images were acquired with an inverted widefield fluorescence microscope Axio Observer 7 (ZEISS, Germany) using appropriate filters and a 10x EC Plan-NEOFLUAR objective or a 1.3 NA 100x EC Plan-NEOFLUAR oil objective in combination with an AxioCam MRm CCD camera. Exposure times were defined empirically and differed depending on the probe sets and fluorescent dyes. For the FISH-immunofluorescence assay, Z-stacked images of infected cells and immobilized virions were acquired with a fixed interval of 0.28-0.31 μm between slices. Raw images were deconvolved in standard mode using Huygens Professional version 21.04 (Scientific Volume Imaging B.V., The Netherlands). If required, raw images were Z-aligned in ZEN 2.6 Pro (ZEISS, Germany) before deconvolution. For analysis, 3D data were converted to maximum intensity projections using Z-project within ImageJ ^39^. Detection, quantification and colocalization analyses of individual spots, each representing a single virion or vRNP, were performed in ImageJ in combination with the plugin ComDet version 0.5.0 (https://github.com/ekatrukha/ComDet). Spot detection thresholds for each channel were set empirically by individual examination of images. The threshold to define co-localized spots was set to a maximum distance of 3-4 pixels between the centers of the spots. For visualization purposes, image brightness and contrast were manually adjusted in ImageJ.

### Mosquito experiment

Adult *Aedes aegypti* (Rockefeller strain) mosquitoes were reared in 30 × 30 × 30 cm cubic cages at 27 ± 1°C and a 12:12 h light:dark cycle, at the Laboratory of Entomology, Wageningen University & Research. Five groups of mosquitoes, housed independently in small cardboard buckets, were allowed to feed through a Parafilm M membrane for at least 1.5 h on 37°C-heated virus-spiked blood meals using a Hemotek PS5 membrane feeding system (Hemotek Ltd, United Kingdom). Virus-spiked blood meals were prepared by mixing virus stocks with bovine blood washed twice with DPBS in a 2:1 ratio (**Table 1**). Virus-blood mixtures were back-titrated as described above. After feeding, mosquitoes were anesthetized using a semi-permeable CO_2_ pad. Engorged female mosquitoes were selected, placed into new cardboard buckets and maintained at 28°C. Throughout the duration of the experiment, mosquitoes were provided with a cotton pad soaked in a 6% sucrose solution *ad libitum*, except for the day before feeding and the day before forced salivation, in which mosquitoes were provided only with water *ad libitum*. To evaluate whether feeding on a blood meal spiked with a mixture of incomplete virus populations can result in virus infection and transmission by mosquitoes, body and saliva samples were collected and tested by virus isolation. Briefly, 12-15 days after the blood meal, mosquitoes were anesthetized using a semi-permeable CO_2_ pad, wings and legs were removed with forceps, and mosquitoes were forced to salivate by inserting their proboscis inside a 20 μL filter tip pre-filled with 7 μL of a 1:1 FBS-50% sucrose mixture for at least 1 h. After salivation, mosquito bodies were collected and stored at -80°C until further processing. Data corresponding to mosquitoes from groups #1, #2 and #3 derive from a single biological experiment. Data corresponding to mosquitoes from groups #4 and #5 derive from two independent experiments.

### Virus isolation

Saliva samples were directly added onto 3 × 10^4^ BSR-T7/5 cells/well previously seeded in 96-well plates (groups #1, #2 and #3) or used to prepare four serial ten-fold dilutions, starting at 1:10 (groups #4 and #5). In the latter case, 50 μL of each dilution were added onto 3 × 10^4^ BSR-T7/5 cells/well previously seeded in 96-well plates. Mosquito bodies were suspended in 100 μL of complete G-MEM additionally supplemented with an extra 1% antibiotic/antimycotic, homogenized using a pellet pestle and handheld pellet pestle motor (Daigger Scientific Inc, USA), and clarified by centrifugation at maximum speed. The supernatants were diluted with additional 200 μL of supplemented G-MEM and used to prepare eight serial dilutions (five-fold the first dilution and ten-fold the remaining dilutions). 50 μL of each dilution were added onto 3 × 10^4^ BSR-T7/5 cells/well previously seeded in 96-well plates. For both saliva and body samples, the expression of the respective fluorescent protein in plaques of infected cells was monitored by fluorescence microscopy 24-72 h post-infection and used as the read-out method.

### Modeling the fraction of infected and co-infected cells

To model the relationship between MOI and the fraction of (co-)infected cells, we considered predictions of three models. Here we provide a brief summary. A complete description of the model and the code are provided as **Supplementary File 1**. Model A predicts the fraction of infected and co-infected cells based on information on the MOI and the frequency of the two variants (iRVFV-SL-eGFP and iRVFV-SL-mCherry2), following a previously described model based on the Poisson distribution^40^. This model has no free parameters to fit, and assumes each virus particle is infectious and complete. Model B introduces non-selective packaging of genome segments, assuming a random distribution of the segment types (in the absence of detailed information for this setup). This model has one free parameter, the probability of infection *p*, in effect allowing the realized MOI to deviate from that set experimentally. Model C introduces variation in the susceptibility of cells, which follows a *β*-distribution over cells. These differences in susceptibility are likely to arise for many reasons, *e*.*g*. differences in the cell growth phase. This model has three parameters: the infection probability *p*, and the two shape parameters for the distribution of susceptibility *α* and *β*. Model C led to a much-improved fit (see **Supplementary File 1**) and hence was selected for visualization.

### Modeling virus spread and the relationship between MOI and co-infection

We generated a simple simulation model of viral spread to consider the impact of non-selective packaging and infection by incomplete virus particles on within-host dynamics. Here we provide a brief summary. A complete description of the model and the code are provided as **Supplementary File 2**. We modeled virus expansion in a fixed number of cells, over discrete viral generations and such that the virus particles released can only infect cells in the next round of infection. We assumed that each infected cell produces a fixed number of virus particles, and that the number of virus particles entering cells follows a Poisson distribution (the mean of this distribution being the MOI). We considered the distribution of the three genome segments over these virus particles. We assumed there is a productive infection only when all three genome segments are present in a cell. However, as described in the results section, we varied the rules governing the transmission between cells (only by complete virus particles, or by both complete and incomplete virus particles) and the distribution of genome segments over virus particles (non-selective, following approximately the empirical distribution, or selective). As the selective packager does not produce incomplete virus particles, this results in three scenarios. For the selective packager, we assumed the total number of virus particles that could be generated was limited by the number of virus genome segments in the infected cell, resulting in the production of a smaller number of virus particles (but containing the same total number of genome segments as the non-selective packager). We then allowed the number of infected cells to expand until all cells became infected or the virus population went extinct.

To consider the relationship between MOI and the fraction of cells infected only by incomplete particles, we first considered the trends for the simulations in a fixed number of cells described above. However, if we fix the mean MOI, we can consider model predictions systematically over a wide range of MOI values. To test the generality of the observations obtained, we randomly drew values for the frequency of each segment type from a uniform distribution, and then normalized these values by the sum of all drawn values (**Supplementary File 2**).

## Data analysis and visualization

Model predictions were performed and plotted in R version 4.0.4 (https://www.R-project.org/). Prism 9 (GraphPad Software) was used to generate graphs of all the remaining results. Sample size varied per experiment and is indicated in each figure legend.

## Acknowledgements

E.B.M is a grateful recipient of scholarships from the Graduate School for Production Ecology & Resource Conservation (PE&RC) and Universidad de Costa Rica (OAICE-031-2019). We thank Michèle Bouloy (Institut Pasteur, France), Karl-Klaus Conzelmann (Ludwig-Maximilians-Universität München) and Connie Schmaljohn (US Army Medical Research Institute of Infectious Diseases) for previously providing the RVFV strain Clone 13, the BSR-T7/5 cells and the antibody 4-D4, respectively. We also thank Lars Ravesloot (Wageningen Bioveterinary Research) and Marcel H. Tempelaars (Shared Research Facilities, Wageningen University & Research) for technical assistance in flow cytometry. Illustrations in Figs. 1a, 1d, 2a, 2d, 3a, 4a, 5a, 6a, 8a and 8b were created with BioRender.com.

## Author contributions

E.B.M, P.J.W.S and J.K conceived the project. E.B.M, P.J.W.S and K.F.B designed the experiments. E.B.M and K.F.B performed the reverse genetics, infection and immunofluorescence experiments. E.B.M performed the growth curves, flow cytometry and smFISH-immunofluorescence experiments. K.F.B performed the IPMA and RT-qPCR experiments. C.J.M.K provided the mosquitoes and bovine blood. E.B.M, K.F.B, S.v.d.W, I.C.R, R.P.M.V and P.J.W.S performed the mosquito experiments. M.P.Z. performed the mathematical modeling with contributions of E.B.M, J.K and P.J.W.S. E.B.M, K.F.B, M.P.Z and P.J.W.S analyzed and interpreted the data with contributions of G.P.P. and J.K. P.J.W.S, J.K. and G.P.P supervised the project. E.B.M wrote the manuscript with contributions of M.P.Z and P.J.W.S. E.B.M and M.P.Z. made the figures with contributions of K.F.B and P.J.W.S. All authors reviewed the manuscript and provided feedback.

## References

1. Michalakis, Y. & Blanc, S. The Curious Strategy of Multipartite Viruses. Annual Review of Virology 7, 203–218 (2020).

2. Sicard, A. et al. Gene copy number is differentially regulated in a multipartite virus. Nature Communications 4, 1–8 (2013).

3. Sicard, A. et al. A multicellular way of life for a multipartite virus. eLife 8, e43599 (2019).

4. Sicard, A., Michalakis, Y., Gutiérrez, S. & Blanc, S. The Strange Lifestyle of Multipartite Viruses. PLoS Pathogens 12, e1005819 (2016).

5. Lucía-Sanz, A. & Manrubia, S. Multipartite viruses: adaptive trick or evolutionary treat? npj Systems Biology and Applications 3, 1–11 (2017).

6. Noda, T. et al. Architecture of ribonucleoprotein complexes in influenza A virus particles. Nature 439, 490–492 (2006).

7. Noda, T. et al. Three-dimensional analysis of ribonucleoprotein complexes in influenza A virus. Nature Communications 3, 1–6 (2012).

8. Chou, Y. et al. One influenza virus particle packages eight unique viral RNAs as shown by FISH analysis. PNAS 109, 9101–9106 (2012).

9. Fournier, E. et al. A supramolecular assembly formed by influenza A virus genomic RNA segments. Nucleic Acids Res 40, 2197–2209 (2012).

10. Goto, H., Muramoto, Y., Noda, T. & Kawaoka, Y. The Genome-Packaging Signal of the Influenza A Virus Genome Comprises a Genome Incorporation Signal and a Genome-Bundling Signal. Journal of Virology 87, 11316–11322 (2013).

11. Dadonaite, B. et al. The structure of the influenza A virus genome. Nature Microbiology 4, 1781–1789 (2019).

12. Le Sage, V. et al. Mapping of Influenza Virus RNA-RNA Interactions Reveals a Flexible Network. Cell Reports 31, (2020).

13. Wichgers Schreur, P. J., Kormelink, R. & Kortekaas, J. Genome packaging of the Bunyavirales. Current Opinion in Virology 33, 151–155 (2018).

14. Wichgers Schreur, P. J. & Kortekaas, J. Single-Molecule FISH Reveals Non-selective Packaging of Rift Valley Fever Virus Genome Segments. PLoS Pathogens 12, e1005800 (2016).

15. Bermúdez-Méndez, E., Katrukha, E. A., Spruit, C. M., Kortekaas, J. & Wichgers Schreur, P. J. Visualizing the ribonucleoprotein content of single bunyavirus virions reveals more efficient genome packaging in the arthropod host. Communications Biology 4, 1–13 (2021).

16. Kortekaas, J. et al. Creation of a Nonspreading Rift Valley Fever Virus. Journal of Virology 85, 12622–12630 (2011).

17. Manzoni, T. B. & López, C. B. Defective (interfering) viral genomes re-explored: impact on antiviral immunity and virus persistence. Future Virology 13, 493–503 (2018).

18. Bara, J. J. & Muturi, E. J. Effect of mixed infections of Sindbis and La Crosse viruses on replication of each virus in vitro. Acta Tropica 130, 71–75 (2014).

19. Pereira Abrao, E. & Lopes da Fonseca, B.A. Infection of Mosquito Cells (C6/36) by Dengue-2 Virus Interferes with Subsequent Infection by Yellow Fever Virus. Vector-Borne and Zoonotic Diseases 16, 124–130 (2016).

20. Wernike, K., Brocchi, E. & Beer, M. Effective interference between Simbu serogroup orthobunyaviruses in mammalian cells. Veterinary Microbiology 196, 23–26 (2016).

21. Boussier, J. et al. Chikungunya virus superinfection exclusion is mediated by a block in viral replication and does not rely on non-structural protein 2. PLoS ONE 15, e0241592 (2020).

22. Jacobs, N. T. et al. Incomplete influenza A virus genomes occur frequently but are readily complemented during localized viral spread. Nature Communications 10, 1–17 (2019).

23. Olson, K. E. et al. Development of a Sindbis virus expression system that efficiently expresses green fluorescent protein in midguts of Aedes aegypti following per os infection. Insect Molecular Biology 9, 57–65 (2000).

24. Scholle, F., Girard, Y. A., Zhao, Q., Higgs, S. & Mason, P. W. trans-Packaged West Nile Virus-Like Particles: Infectious Properties In Vitro and in Infected Mosquito Vectors. Journal of Virology 78, 11605–11614 (2004).

25. Foy, B. D. et al. Development of a new Sindbis virus transducing system and its characterization in three Culicine mosquitoes and two Lepidopteran species. Insect Molecular Biology 13, 89–100 (2004).

26. Smith, D. R., Adams, A. P., Kenney, J. L., Wang, E. & Weaver, S. C. Venezuelan equine encephalitis virus in the mosquito vector Aedes taeniorhynchus: Infection initiated by a small number of susceptible epithelial cells and a population bottleneck. Virology 372, 176–186 (2008).

27. Weaver, S. C., Forrester, N. L., Liu, J. & Vasilakis, N. Population bottlenecks and founder effects: implications for mosquito-borne arboviral emergence. Nature Reviews Microbiology 19, 184–195 (2021).

28. Cuevas, J. M., Durán-Moreno, M. & Sanjuán, R. Multi-virion infectious units arise from free viral particles in an enveloped virus. Nat Microbiol 2, 1–7 (2017).

29. Sanjuán, R. Collective Infectious Units in Viruses. Trends in Microbiology 25, 402–412 (2017).

30. Andreu-Moreno, I. & Sanjuán, R. Collective Infection of Cells by Viral Aggregates Promotes Early Viral Proliferation and Reveals a Cellular-Level Allee Effect. Current Biology 28, 3212-3219.e4 (2018).

31. Sanjuán, R. & Thoulouze, M.-I. Why viruses sometimes disperse in groups? Virus Evolution 5, (2019).

32. Buchholz, U. J., Finke, S. & Conzelmann, K.-K. Generation of Bovine Respiratory Syncytial Virus (BRSV) from cDNA: BRSV NS2 Is Not Essential for Virus Replication in Tissue Culture, and the Human RSV Leader Region Acts as a Functional BRSV Genome Promoter. Journal of Virology 73, 251–259 (1999).

33. Muller, R. et al. Characterization of clone 13, a naturally attenuated avirulent isolate of Rift Valley fever virus, which is altered in the small segment. Am. J. Trop. Med. Hyg. 53, 405–411 (1995).

34. Barnard B.J.H. Rift Valley fever vaccine -antibody and immune response in cattle to a live and an inactivated vaccine. Journal of the South African Veterinary Association 50, 155–157 (1979).

35. Femino, A. M., Fay, F. S., Fogarty, K. & Singer, R. H. Visualization of Single RNA Transcripts in Situ. Science 280, 585–590 (1998).

36. Raj, A., van den Bogaard, P., Rifkin, S. A., van Oudenaarden, A. & Tyagi, S. Imaging individual mRNA molecules using multiple singly labeled probes. Nature Methods 5, 877–879 (2008).

37. Orjalo Jr., A., Johansson, H. E. & Ruth, J. L. StellarisTM fluorescence in situ hybridization (FISH) probes: a powerful tool for mRNA detection. Nature Methods 8, 884 (2011).

38. Keegan, K. & Collett, M. S. Use of bacterial expression cloning to define the amino acid sequences of antigenic determinants on the G2 glycoprotein of Rift Valley fever virus. Journal of Virology 58, 263–270 (1986).

39. Schneider, C. A., Rasband, W. S. & Eliceiri, K. W. NIH Image to ImageJ: 25 years of image analysis. Nature Methods 9, 671–675 (2012).

40. Zwart, M. P. et al. An experimental test of the independent action hypothesis in virus–insect pathosystems. Proceedings of the Royal Society B: Biological Sciences 276, 2233–2242 (2009).

