## Supplementary Information for "Incomplete bunyavirus particles contribute to within-host spread and between-host transmission"

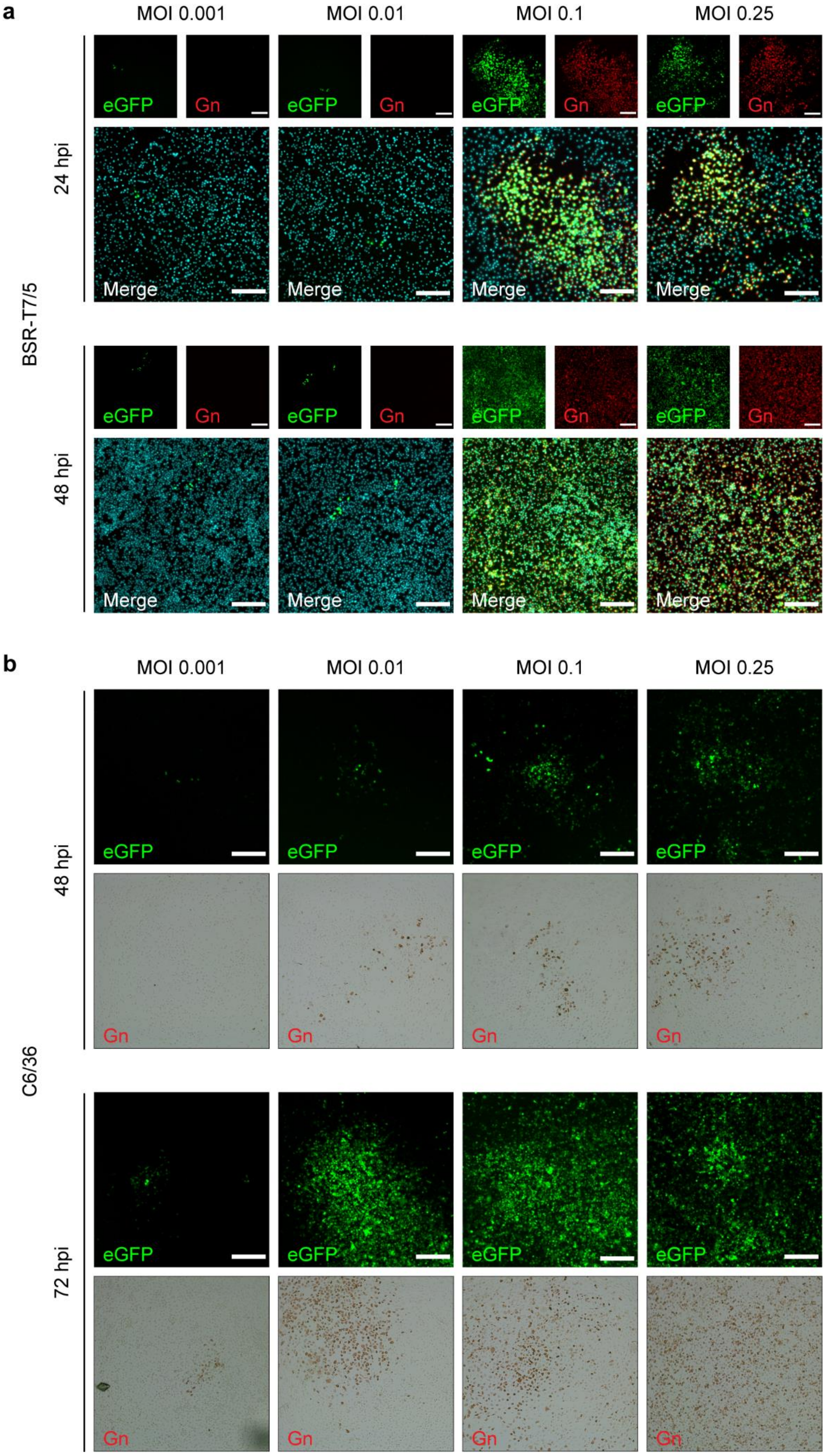
a
MOI 0.001
MOI 0.01
MOI 0.1
MOI 0.25
BSR-T7/5
24 hpi
48 hpi
eGFP
Gn
Merge
b
MOI 0.001
MOI 0.01
MOI 0.1
MOI 0.25
C6/36
48 hpi
72 hpi
eGFP
Gn

**Supplementary Fig. 1 Co-infection of mammalian and insect cells with complementing incomplete RVFV particles. a, b** Mammalian (BSR-T7/5) (**a**) and insect (C6/36) (**b**) cells were simultaneously infected with non-spreading iRVFV-SL-eGFP and iRVFV-ML particles at increasing MOIs (ranging from 0.001 to 0.25 for each virus). Co-infection with the two populations of incomplete RVFV particles supports genome complementation, allows virus replication, production of infectious progeny and virus spread. Infected cells were analyzed at 24-48 h (BSR-T7/5 cells) or 48-72 h (C6/36 cells) post-infection by following the expression of eGFP (green) via direct fluorescence microscopy examination and the expression of Gn (red) via an immunofluorescence assay in BSR-T7/5 cells or an immunoperoxidase monolayer assay in C6/36 cells. Expression of Gn was detected with rabbit polyclonal anti-Gn serum in combination with Alexa Fluor 568-conjugated secondary antibodies (immunofluorescence assay) or with HRP-conjugated secondary antibodies (immunoperoxidase monolayer assay). Cell nuclei (cyan) were visualized with DAPI. Of note, with the sole intention of depicting the outcome progression at increasing MOIs, images corresponding to co-infections (MOI of 0.1) at 24 h (BSR-T/5) and 72 h (C6/36) post-infection were purposely selected to be the exact same images as shown in **Fig. 4**. Scale bars, 200 µm.

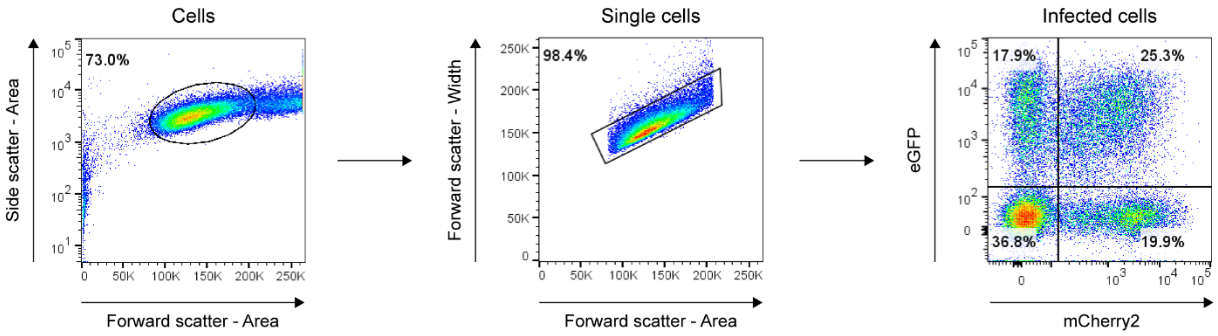


**Supplementary Fig. 2 Flow cytometry gating strategy.** Illustrative example of the gating strategy employed for the analysis of flow cytometry data. BSR-T7/5 cells were mock-infected, singly-infected or co-infected with non-spreading iRVFV-SL-eGFP and/or iRVFV-SL-mCherry2 particles. The cell population of interest was first discriminated from debris. Then, a gate was applied to select single cell events from doublets. Finally, we quantified the fraction of non-infected, singly-infected or co-infected cells by determining the expression of eGFP, mCherry2 or both. The plots depicted here correspond to the co-infected sample (MOI of 0.5 for each virus population), as shown in **Fig. 2**.

**Supplementary Table 1** Primers for cDNA synthesis of viral genome segments.

| Target | Name | Sequence |
|---|---|---|
| RVFV-Clone 13-S and RVFV-35/74-S | JR860-For | ACAAAGCTCCCTAGAGATACA |
| RVFV-Clone 13-M and RVFV-35/74-M | JR861-For | GACACAAAGACGGTGCATTA |
| RVFV-Clone 13-L and RVFV-35/74-L | JR890-For | GACACAAAGGCGCCCAATC |

**Supplementary Table 2** Primers for RT-qPCR amplifications of viral genome fragments and *ampR*.

| Target | Name | Sequence |
|---|---|---|
| RVFV-Clone 13-S and RVFV-35/74-S | JR907-For | TCCAGTTTGCTGCTCAA |
| | JR908-Rev | CTGCTTTAAGAGTTCGATAACC |
| RVFV-Clone 13-M and RVFV-35/74-M | JR909-For | GCTGATGGCTTGAACAAC |
| | JR910-Rev | GTCTCTCACACCGAACTATC |
| RVFV-Clone 13-L and RVFV-35/74-L | JR911-For | TCGATAGATGTGGAAGATATGG |
| | JR912-Rev | CGTCATTCATCATGGGAAAC |
| *ampR* | JR971-For | GCAGTGTTATCACTCATGG |
| | JR972-Rev | CACTATTCTCAGAATGACTTGG |

**Supplementary Table 3** Antibodies used in immunostaining assays.

| Assay | Target | Antibody | Dilution | Source/reference |
|---|---|---|---|---|
| IPMA, IF (1$^{ary}$) | RVFV Gn | Rabbit polyclonal serum | 1:500 | Thermo Fisher |
| IPMA (2$^{ary}$) | Rabbit IgG | Goat polyclonal anti-rabbit IgG HRP-conjugated | 1:500 | P0448 Dako |
| IF (2$^{ary}$) | Rabbit IgG | Goat polyclonal anti-rabbit IgG-FITC | 1:250 | sc-2012 Santa Cruz Biotechnology |
| IF (2$^{ary}$) | Rabbit IgG | Donkey polyclonal anti-rabbit IgG-Alexa Fluor 568 | 1:500 | A10042 Invitrogen |
| IF (1$^{ary}$) | RVFV N | Monoclonal mouse hybridoma | 1:100 | F1D11 CISA-INIA |
| IF (2$^{ary}$) | Mouse IgG | Goat polyclonal anti-mouse IgG-Alexa Fluor Plus 488 | 1:500 | A32723 Invitrogen |
| FISH-IF (1$^{ary}$) | RVFV Gn | Hybridoma 4-D4 supernatant | 1:160 | [1] |
| FISH-IF (2$^{ary}$) | Mouse IgG | Goat polyclonal anti-mouse IgG-Alexa Fluor 488 | 1:1000 | A-11001 Invitrogen |

IPMA: immunoperoxidase monolayer assay, IF: immunofluorescence, FISH-IF: fluorescence *in situ* hybridization-immunofluorescence.

### Supplementary References


1. Keegan, K. & Collett, M. S. Use of bacterial expression cloning to define the amino acid sequences of antigenic determinants on the G2 glycoprotein of Rift Valley fever virus. *Journal of Virology* **58**, 263–270 (1986).
