## Supplementary File 1 for "Incomplete bunyavirus particles contribute to within-host spread and between-host transmission": Supplementary File 1.html

Modeling the fraction of infected and co-infected cells


### Modeling the fraction of infected and co-infected cells

###### Author: Mark P. Zwart

###### Date: 14 February 2022

##### Supplementary File 1

###### Manuscript: “Incomplete bunyavirus particles contribute to within-host spread and between-host transmission”

###### Erick Bermúdez-Méndez, Kirsten F. Bronsvoort, Mark P. Zwart, Sandra van de Water, Ingrid Cárdenas-Rey, Rianka P. M. Vloet, Constantianus J. M. Koenraadt, Gorben P. Pijlman, Jeroen Kortekaas, Paul J. Wichgers Schreur

###### Intro

Experimental data set: hamster-derived BSR-T7/5 cells were infected at increasing values of cellular multiplicity of infection (MOI) with two RVFV variants labeled with fluorescent markers (Fig. 2 in the manuscript). Flow cytometry was then used to determine how many cells were co-infected with the two different virus variants. The key question is whether the rate of cellular co-infection (as determined by yellow fluorescence, indicating both red and green fluorescent proteins are present), can be predicted and is compatible with theoretical predictions. We will attempt to predict the rate of cellular co-infection from the rate of cellular infection with a simple model (Model A), and fit more complex models that predict the fraction of cells infected by none virus variants, one of the two variants, or both variants (Models B and C).

###### Experimental data

First, we load the experimental data. These are flow cytometry-derived cell counts, coming from a single experiment. In the model fitting, we treat each cell as an independent observation, whereas in reality this is only technical and not biological replication (i.e. in real life the experimental variation will be larger than assumed in the model fitting described here). The indicator of model fit (i.e. the negative log likelihood) must therefore be interpreted cautiously, and only very obvious differences in model fit are likely to be meaningful. We make a new array with only the data that are informative here, removing the controls in which none or only one virus is included in the inoculum.

```
# Load the data from a *.csv file.
setwd("C:/R_Output_EBM/Manuscript (Incomplete RVFV particles)")
c.data <- read.csv(file = "Data_Supplementary File 1_v1.csv", header = TRUE)
print(c.data)
```

```
##   MOI_eGFP MOI_mCherry2 Uninf_cells eGFP_cells mCherry2_cells Both_marker_cells
## 1     0.00         0.00       32716          0              0                 0
## 2     0.50         0.00       23231      13546              0                 0
## 3     0.00         0.50       20059          0          17276                 0
## 4     0.10         0.10       27376       3162           3402              1415
## 5     0.25         0.25       23149       4909           6367              3267
## 6     0.50         0.50       13232       6439           7147              9095
## 7     1.00         1.00        7895       6731           7554             16034
## 8     2.50         2.50        2749       5160           5306             25234
## 9       NA           NA          NA         NA             NA                NA
```

```
# New array without controls that are not meaningful for this analysis, and 
# calculate some numbers that will be useful later in different analyses.
t.data <- c.data[4:8,]
moi.vals = rowSums(t.data[,1:2])
all.cells = rowSums(t.data[,3:6])
inf.cells = rowSums(t.data[,4:6])

# Determine whether there is a bias towards one variant in the experiment.
ino.gfp = c.data[2,4]/sum(c.data[2,3:6])
ino.che = c.data[3,5]/sum(c.data[3,3:6])
p.f.gfp = ino.gfp/(ino.gfp+ino.che)
```

The more complex Models B and C include various mechanisms that may contribute to an improved model fit. These models are all inspired by those in an unpublished manuscript with Lia Hemerik and Wopke van der Werf on infection kinetics. To fit these models with existing code, first we reformat the data. Note that the names used in this code (infected and dead) reflect that it is used for describing infection of animals and plants by pathogenic bacterial and viral strains.

```
doses <- moi.vals
s.uninfected <- t.data[,3]
s.infected.a <- t.data[,4]
s.infected.b <- t.data[,5]
s.infected.ab <- t.data[,6]
s.dead <- inf.cells
s.total <- all.cells
num.doses <- length(doses)
inoculum.freq <- p.f.gfp
```

###### Model A: Simple MOI-derived prediction

First, we plot the fraction of infected (any RVFV variant) and co-infected (both RVFV variants) cells to start off with. We will also include the simplest and most direct approach for predicting cellular infection and co-infection: to do so directly from the MOI. For a complete explanation see for example Zwart et al. 2013 (Model-Selection-Based Approach for Calculating Cellular Multiplicity of Infection during Virus Colonization of Multi-Cellular Hosts. PLoS ONE 8(5): e64657. https://doi.org/10.1371/journal.pone.0064657). In essence, we assume the number of infecting virus particles over cells follows a Poisson distribution, use the fraction of uninfected cells to predict the mean of the Poisson distribution, and include information on the frequency of the two variants to predict the rate of co-infection.

```
# Fraction of infected and co-infected cells
f.inf = inf.cells/all.cells
f.coinf = t.data[,6]/all.cells

# Plot of the data
par(mar = c(5, 6, 4, 2))
plot(x = log2(moi.vals), y = f.inf, cex = 1.5,
     main = "Model A",  cex.main = 2, font.main = 1,
     xlab = "", ylab = "", 
     type = "p", col = "blue", pch = 19, 
     xlim = c(-4, 4), ylim = c(0, 1),
     cex.lab = 2, cex.axis = 2,
     las = 1)
points(x = log2(moi.vals), y = f.coinf, type = "p", col = "magenta", pch = 19, cex = 1.5)
title(xlab = expression("log"[2]*" MOI"), cex.lab = 2)
title(ylab = expression("Infected and co-infected cells"), line = 3.7, cex.lab = 2)

# Predict and plot the fraction of infected cells from inoculum MOI
hyp.moi.vals = 2^seq(-4, 4, by = 0.01)
pred.inf = 1 - exp(-hyp.moi.vals)
lines(x = log2(hyp.moi.vals), y = pred.inf, type = "l", col = "blue", lty = 2, lwd = 3)

# Predict and plot the fraction of co-infected cells from inoculum MOI. 
pred.coinf = (1 - exp(-p.f.gfp*hyp.moi.vals))*(1 - exp(-(1-p.f.gfp)*hyp.moi.vals))
lines(x = log2(hyp.moi.vals), y = pred.coinf, type = "l", col = "magenta", lwd = 3)
```

When we graph the experimental data and predictions from the simplest of the models, we see that the data are surprisingly congruent with model predictions. In the figure above, the \(log\_2\) of MOI is given on the x axis, and the response is given on the y axis. In this and in subsequent plots, the fraction of infected cells experimental data are indicated with blue circles, and the model predictions with the dashed blue line. The fraction of co-infected cells experimental data are indicated with magenta circles, and the model predictions with the solid magenta line. Clearly, there is some discrepancy between the model and the data, as the observed response for both infection and co-infection appear to be more gradual than the simple model predictions. We can use a modeling framework to try and better understand these patterns, and what mechanisms may account for these discrepancies.

To be able to compare Model A to Models B and C, we calculate the negative log likelihood as we will do subsequently for the other two models. However, as this model prediction uses a completely different approach to the other two models and has no free parameters to be fitted, this value is only calculated to give a rough indication of model fit.

```
# To get some indication of how well this model fits the data and to compare
# this model to the more complex models, we can calculate the log likelihood.
# This calculation is done using the same setup as done below for Models B and C.

# Make predictions of the rate of non-infected cells, co-infected cells, and
# cells infected with only one variant.
pred.none = exp(-moi.vals)
pred.ab = (1 - exp(-p.f.gfp*moi.vals))*(1 - exp(-(1-p.f.gfp)*moi.vals))
pred.a  = (1 - exp(-p.f.gfp*moi.vals))*exp(-(1-p.f.gfp)*moi.vals)
pred.b  = exp(-p.f.gfp*moi.vals)*(1 - exp(-(1-p.f.gfp)*moi.vals))

# Next calculate the log likelihoods for each dose.
mort <- array(NA, dim=c(nrow=0,ncol=1))
    
for (i in 1:num.doses) {
    
  # Calculate likelihood of the predicted mortality.
    data.for.comp <- c(s.uninfected[i], s.infected.a[i], s.infected.b[i], s.infected.ab[i])
    model.for.comp <- c(pred.none[i], pred.a[i], pred.b[i], pred.ab[i])
    likelihood = dmultinom(x = data.for.comp, size = NULL, prob = model.for.comp, log = TRUE)
                
    # Send to output array 
    mort=rbind(mort, likelihood)

}   # end i loop

# Determine the NLL over all doses.
new.nll = -sum(mort[,1])
print(new.nll)
```

```
## [1] 10954.69
```

###### Model B: Non-selective packaging

Model A gives a surprisingly good account of the data given its simplicity and lack of any free parameters to be fitted. On the other hand, there appear to be discrepancies between the model predictions and the data. We therefore consider more complex models that might account for these discrepancies. First, we will consider the importance of the assumption that virus genome segments are randomly packaged into the virus particles used to generate the inoculum. Since we do not have information on what the empirical distribution of segments per virus particle looks like for these experiments, we will assume that each virus particle packages three segments, and that the 2 segments present in the replicon-infected cell population occur at equal frequencies and are then packaged at random into the particles.

```
gf.a = 0.5  # the "genome formula": the relative frequency of segment 1
gf.b = 0.5
seg.per.virpar = 3
```

First, we set the parameter space in which we can pick the initial value of the parameters to be fit. For Model 1 this is a single parameter \(p\), the probability of infection. Note that the range only constricts the parameter space in which the search can start, not where the values the parameter can take during the search. Since Model A gives a reasonable prediction, and based on some preliminary model fitting runs, we can take a narrow starting parameter space as we expect values close to one for the parameter \(p\).

```
# Number of parameters to be estimated
model.parameters = 1

# p, probablity of infection
log.p.unit = 0.01
log.p.range <- seq(-0.5, 0.5, by = log.p.unit)
```

Now set the remaining parameters for the search, and generate an array to store the final data produced during the searches. Note that the models without HHS (see below) could easily be predicted analytically instead of iteratively, but to keep everything comparable we use an iterative approach throughout model fitting. To ensure these results are reproducible, we also set the random seed here.

```
# Finally set the limitations on the search
max.iter = 5000      # max number of total iterations
max.iter.stuck = 20  # max number iterations without improvement
repititions = 100    # the number of times the complete search is performed                    
simulations = 10^5   # the number of individual hosts for which the infection 
                     # process is iterated to get a model prediction. 
lambda.limit = 10000 # Limit the lambda value to avoid numerical overflow 
                     # issues. Should be set to a high value, so you are not 
                     # drawing any zero values from the Poisson distribution
                     # with mean lambda.

# Generate an array to store final data including model parameter estimates.
final.data <- array(NA, dim=c(0,4))
colnames(final.data) <- c("Iterations", "Stuck_Counter", "log.p", "NLL")

# For reproducibility, set the random seed
set.seed(42)
```

Next we run the loop for the actual stochastic hill climbing search. Note that because this is a generic algorithm and because it relies on a high number of iterations to obtain a model prediction, it is not exactly fast.

```
# Then the SHC loop itself

for(h in 1:repititions) {
    
    # Randomly pick start values for the search
    log.p = sample(log.p.range, size = 1)
        
    # Housekeeping: set iterations to 1, exit.flag indicates when to end the search, and set
    # NLL to a very high value so that the first value obtained is accepted.
    iter = 1
    exit.flag = 0
    nll = 1e15
    stuck.counter = 0

    while(exit.flag == 0) {
        
        # Choose model parameter to mutate and then mutate it, antilog transform all model param.
        choose.param = sample(1:model.parameters, size = 1)
        mutate.up.or.down = sample(c(1, -1), size = 1)
        if(choose.param == 1) new.log.p = log.p + (log.p.unit*mutate.up.or.down) else new.log.p = log.p
        p = 10^new.log.p
        
        # Now evaluate the model for these parameters and determine the NLL. Here is
        # a loop for which you do this for every dose. The array mort stores this 
        # information for each dose.
            
        mort <- array(0:0, dim=c(nrow=0,ncol=5))
    
        for (i in 1:num.doses) {
            
            something = rep( (p*doses[i]), simulations)
            lambda.a = something*inoculum.freq  # Here the code is modified to limit size of lambda values.
            big.a = which(lambda.a > lambda.limit)
            lambda.a[big.a] = lambda.limit  
            lambda.b = something*(1-inoculum.freq)
            big.b = which(lambda.b > lambda.limit)
            lambda.b[big.b] = lambda.limit              
            infectors.a = rpois(n = simulations, lambda = lambda.a)
            infectors.b = rpois(n = simulations, lambda = lambda.b)
            infectors.a.1 = rbinom(n = simulations, size = (seg.per.virpar*infectors.a), prob = gf.a)       
            infectors.a.2 = (seg.per.virpar*infectors.a) - infectors.a.1
            infectors.b.1 = rbinom(n = simulations, size = (seg.per.virpar*infectors.b), prob = gf.b)
            infectors.b.2 = (seg.per.virpar*infectors.b) - infectors.b.1
            infectors.1 = infectors.a.1 + infectors.b.1 
            infectors.2 = infectors.a.2 + infectors.b.2 
            infected = which(infectors.1 > 0 & infectors.2) 
            survival = (simulations - length(infected))/simulations
            infection.ab = length(which(infectors.a.2[infected] > 0 & infectors.b.2[infected] > 0))/simulations
            infection.a = length(which(infectors.a.2[infected] > 0 & infectors.b.2[infected] == 0))/simulations
            infection.b = length(which(infectors.a.2[infected] == 0 & infectors.b.2[infected] > 0))/simulations
            
            # For some reason very small negative values pop up
            # sometimes, hence the insurance below.
            if(survival < 0) survival = 0       
            if(infection.a < 0) infection.a = 0 
            if(infection.b < 0) infection.b = 0
            if(infection.ab < 0) infection.ab = 0
                
            # Calculate likelihood of the predicted mortality.
            data.for.comp <- c(s.uninfected[i], s.infected.a[i], s.infected.b[i], s.infected.ab[i])
            model.for.comp <- c(survival, infection.a, infection.b, infection.ab)
            likelihood = dmultinom(x = data.for.comp, size = NULL, prob = model.for.comp, log = TRUE)
                    
            # Send to output array 
            mort=rbind(mort, c(survival, infection.a, infection.b, infection.ab, likelihood))
        
        }   # end i loop
        
        new.nll = -sum(mort[,5])
        
        # Here check if the new parameter values lead to improved fit. If so, accept
        # the new parameter values and NLL, and reset the StuckCounter. Otherwise, 
        # repeat the process with the same values used previously. 
        stuck.counter = stuck.counter + 1
                
        if (new.nll < nll) {
        
            log.p = new.log.p
            nll = new.nll
            stuck.counter = 0
            
            }   # end of if loop
        
        # Check whether to continue the search or exit
        iter = iter + 1
        if (stuck.counter == max.iter.stuck) exit.flag = 1
        if (iter == max.iter) exit.flag = 1 
        
    }  # End of while loop for search  
        
    final.data = rbind(final.data, c(iter, stuck.counter, log.p, nll))
  # print(h)
    
} # end of h loop 

# Write a csv file with the data, to avoid having to repeat this computationally
# intensive step.
setwd("C:/R_Output_EBM/Manuscript (Incomplete RVFV particles)")
write.csv(final.data, file = "Results_Model_B.csv")
```

We order the results obtained and find the lowest NLL value, as this should represent the best set of parameters found. Since we are running multiple independent searches, we print the top ten search results to check if we converged on the best solution multiple times. If not, one might have to run more searches in order to be confident you (i) are running enough searches, and (ii) you are starting in a suitable parameter space for searching (i.e. the solution is within the parameter space and/or the fitting landscape is not too rugged).

```
# Order results based on NLL values
ordered.final.data=order(final.data[,4])
for(j in 1:10) print(final.data[ordered.final.data[j],c(1,3,4)])
```

```
## Iterations      log.p        NLL 
##     49.000      0.030   8587.833 
## Iterations      log.p        NLL 
##     68.000      0.030   8596.464 
## Iterations      log.p        NLL 
##    142.000      0.020   8603.244 
## Iterations      log.p        NLL 
##     29.000      0.020   8605.452 
## Iterations      log.p        NLL 
##     80.000      0.030   8610.629 
## Iterations      log.p        NLL 
##    100.000      0.030   8624.189 
## Iterations      log.p        NLL 
##    118.000      0.030   8630.883 
## Iterations      log.p        NLL 
##     60.000      0.030   8642.505 
## Iterations      log.p        NLL 
##    112.000      0.020   8647.766 
## Iterations      log.p        NLL 
##    156.000      0.020   8648.191
```

Next, to get more confidence about the NLL value, we re-run the model to obtain predictions based on a higher number of iterations. This should give a better approximation of the true NLL value for the set of parameters we have chosen.

```
# Set a higher number of simulations

simulations.check = 1e6

p = 10^final.data[ordered.final.data[1],3]

mort <- array(0:0, dim=c(nrow=0,ncol=5))
    
for (i in 1:num.doses) {
    
    something = rep( (p*doses[i]), simulations.check)
    lambda.a = something*inoculum.freq  
    big.a = which(lambda.a > lambda.limit)
    lambda.a[big.a] = lambda.limit  
    lambda.b = something*(1-inoculum.freq)
    big.b = which(lambda.b > lambda.limit)
    lambda.b[big.b] = lambda.limit              
    infectors.a = rpois(n = simulations.check, lambda = lambda.a)
    infectors.b = rpois(n = simulations.check, lambda = lambda.b)
    infectors.a.or.b = infectors.a + infectors.b
    survival = 1-sum(infectors.a.or.b>0)/simulations.check
    infection.ab = length(which(infectors.a > 0 & infectors.b > 0))/simulations.check
    no.infectors.a = which(infectors.a==0)
    no.infection.a = sum(infectors.a==0)/simulations.check
    infection.a = 1 - no.infection.a - infection.ab
    infection.b = 1 - survival - infection.a - infection.ab 
    
    # For some reason very small negative values pop up
    # sometimes, hence the insurance below.
    if(survival < 0) survival = 0       
    if(infection.a < 0) infection.a = 0 
    if(infection.b < 0) infection.b = 0
    if(infection.ab < 0) infection.ab = 0
            
    # Calculate likelihood of the predicted mortality
    data.for.comp <- c(s.uninfected[i], s.infected.a[i], s.infected.b[i], s.infected.ab[i])
    model.for.comp <- c(survival, infection.a, infection.b, infection.ab)
    likelihood = dmultinom(x = data.for.comp, size = NULL, prob = model.for.comp, log = TRUE)
            
    # Send to output array 
    mort=rbind(mort, c(survival, infection.a, infection.b, infection.ab, likelihood))
    
}   # end i loop

# NLL value
nll.model.1 = -sum(mort[,5])
print(nll.model.1)
```

```
## [1] 12745.99
```

The NLL value obtained for Model B is higher than that for Model A, indicating that this more complex model with a single free parameter does not lead to improved fit. Finally, we plot the data and model predictions, after running the model over a range of doses.

```
# Set an appropriate dose range 
doses.check  = 2^(seq(from = -4, to = 4, by = 0.1))
num.doses.check = length(doses.check)

# Plot the data first
par(mar = c(5, 6, 4, 2))
plot(x = log2(moi.vals), y = f.inf, cex = 1.5,
     main = "Model B", cex.main = 2, font.main = 1,
     xlab = "", ylab = "", 
     type = "p", col = "blue", pch = 19, 
     xlim = c(-4, 4), ylim = c(0, 1),
     cex.lab = 2, cex.axis = 2,
     las = 1)
points(x = log2(moi.vals), y = f.coinf, type = "p", col = "magenta", pch = 19, cex = 1.5)
title(xlab = expression("log"[2]*" MOI"), cex.lab = 2)
title(ylab = expression("Infected and co-infected cells"), line = 3.7, cex.lab = 2)

# Set model parameters
p = 10^final.data[ordered.final.data[1],3]

# make array for results, 
mort <- array(0:0, dim=c(nrow=0,ncol=4))

for (i in 1:num.doses.check) {
    something = rep( (p*doses.check[i]), simulations.check)
    lambda.a = something*inoculum.freq  # Here the code is modified to limit size of lambda values.
    big.a = which(lambda.a > lambda.limit)
    lambda.a[big.a] = lambda.limit  
    lambda.b = something*(1-inoculum.freq)
    big.b = which(lambda.b > lambda.limit)
    lambda.b[big.b] = lambda.limit              
    infectors.a = rpois(n = simulations.check, lambda = lambda.a)
    infectors.b = rpois(n = simulations.check, lambda = lambda.b)
    infectors.a.or.b = infectors.a + infectors.b
    survival = 1-sum(infectors.a.or.b>0)/simulations.check
    infection.ab = length(which(infectors.a > 0 & infectors.b > 0))/simulations.check
    no.infectors.a = which(infectors.a==0)
    no.infection.a = sum(infectors.a==0)/simulations.check
    infection.a = 1 - no.infection.a - infection.ab
    infection.b = 1 - survival - infection.a - infection.ab 

    # No NLL calculation needed here!
    # Send to output array 
    mort=rbind(mort, c(survival, infection.a, infection.b, infection.ab))
        
}   # end i loop
    
# Plot model for this parameter set, infection first
lines(x = log2(doses.check), y = 1-(mort[,1]), col = "blue", lty = 2, lwd = 3)
# Now plot mortality
lines(x = log2(doses.check), y = mort[,4], col = "magenta", lwd = 3)
```

Here we can see that Model B gives a similar prediction to the simple MOI-based model (see above), as was expected from the estimate model parameter values. So this model is not helpful for generating better predictions of cellular co-infection, even though we know that the packaging of viral genome segments is non-selective. We therefore consider other mechanisms that might lead to improved model fit.

###### Model C: Non-selective packaging and heterogeneous host susceptibility

In addition to the Model B assumption on non-selective packaging of genome segments into virus particles, Model C assumes that all hosts, in this case cells, are not equally susceptible to the virus. This is a very plausible assumption, since e.g. depending on the growth phase (G1, S or G2) the susceptibility of cells to viral infection will vary. There could also be other mechanisms that result in heterogeneous susceptibility, such as spatial effects (how exposed is a cell to free virus particles) or possibly genetic variation within the cell population. Following van der Werf et al. 2011 (PLoS Computational Biology 7(6): e1002097. https://doi.org/10.1371/journal.pcbi.1002097), we assume that the probability of infection follows a beta distribution over cells. This flexible distribution allows the model to explore the effects of unimodal or bimodal distributions of host susceptibility. In addition to the probability of infection, the two shape parameters of the beta distribution need to be estimated. Below follows the code for model fitting.

```
# p, probablity of infection: use the same range as for Model A.

# a, scaling parameter of the beta distribution
log.a.unit = 0.01
log.a.range <- seq(-1, 1, by = log.a.unit)

# b, scaling parameter of the beta distribution
log.b.unit = 0.01
log.b.range <- seq(-1, 1, by = log.b.unit)

# Finally alter some limitations on the search
max.iter.stuck = 100  # max number iterations without improvement

# Generate an array to store final data including model parameter estimates.
final.data <- array(NA, dim=c(0,6))
colnames(final.data) <- c("Iterations", "Stuck_Counter", "log.p", "log.a", "log.b", "NLL")
```

We perform the same steps as for Model B above: run the search with a starting parameter space (but now for three free parameters), determine the NLL for a higher number of iterations, and plot the results.

```
for(h in 1:repititions) {
    
    log.p = sample(log.p.range, size = 1)
    log.a = sample(log.a.range, size = 1)
    log.b = sample(log.b.range, size = 1)
    
    iter = 1
    exit.flag = 0
    nll = 1e15
    stuck.counter = 0 

    while(exit.flag == 0) {
        
        choose.param = sample(1:model.parameters, size = 1)
        mutate.up.or.down = sample(c(1, -1), size = 1)
        if(choose.param == 1) new.log.p = log.p + (log.p.unit*mutate.up.or.down) else new.log.p = log.p
        if(choose.param == 2) new.log.a = log.a + (log.a.unit*mutate.up.or.down) else new.log.a = log.a
        if(choose.param == 3) new.log.b = log.b + (log.b.unit*mutate.up.or.down) else new.log.b = log.b
        p = 10^new.log.p
        a = 10^new.log.a
        b = 10^new.log.b

        mort <- array(0:0, dim=c(nrow=0,ncol=5))
    
        for (i in 1:num.doses) {
            
            something = rbeta(n = simulations, shape1 = a, shape2 = b)*p*doses[i]
            lambda.a = something*inoculum.freq  
            big.a = which(lambda.a > lambda.limit)
            lambda.a[big.a] = lambda.limit  
            lambda.b = something*(1-inoculum.freq)
            big.b = which(lambda.b > lambda.limit)
            lambda.b[big.b] = lambda.limit              
            infectors.a = rpois(n = simulations, lambda = lambda.a)
            infectors.b = rpois(n = simulations, lambda = lambda.b)
            infectors.a.1 = rbinom(n = simulations, size = (seg.per.virpar*infectors.a), prob = gf.a)   
            infectors.a.2 = (seg.per.virpar*infectors.a) - infectors.a.1
            infectors.b.1 = rbinom(n = simulations, size = (seg.per.virpar*infectors.b), prob = gf.b)   
            infectors.b.2 = (seg.per.virpar*infectors.b) - infectors.b.1
            infectors.1 = infectors.a.1 + infectors.b.1 
            infectors.2 = infectors.a.2 + infectors.b.2 
            infected = which(infectors.1 > 0 & infectors.2) 
            survival = (simulations - length(infected))/simulations
            infection.ab = length(which(infectors.a.2[infected] > 0 & infectors.b.2[infected] > 0))/simulations 
            infection.a = length(which(infectors.a.2[infected] > 0 & infectors.b.2[infected] == 0))/simulations
            infection.b = length(which(infectors.a.2[infected] == 0 & infectors.b.2[infected] > 0))/simulations

            if(survival < 0) survival = 0       
            if(infection.a < 0) infection.a = 0 
            if(infection.b < 0) infection.b = 0
            if(infection.ab < 0) infection.ab = 0

            data.for.comp <- c(s.uninfected[i], s.infected.a[i], s.infected.b[i], s.infected.ab[i])
            model.for.comp <- c(survival, infection.a, infection.b, infection.ab)
            likelihood = dmultinom(x = data.for.comp, size = NULL, prob = model.for.comp, log = TRUE)

            mort=rbind(mort, c(survival, infection.a, infection.b, infection.ab, likelihood))
        
        }   # end i loop
        
        new.nll = -sum(mort[,5])
        stuck.counter = stuck.counter + 1
                
        if (new.nll < nll) {
        
            log.p = new.log.p
            log.a = new.log.a
            log.b = new.log.b           
            nll = new.nll
            stuck.counter = 0
            
            }   # end of if loop

        iter = iter + 1
        if (stuck.counter == max.iter.stuck) exit.flag = 1
        if (iter == max.iter) exit.flag = 1 
        
    }  # End of while loop for search  
        
    final.data = rbind(final.data, c(iter, stuck.counter, log.p, log.a, log.b, nll))
  # print(h)
    
} # end of h loop 

# Write a csv file with the data, to avoid having to repeat this computationally
# intensive step.
setwd("C:/R_Output_EBM/Manuscript (Incomplete RVFV particles)")
write.csv(final.data, file = "Results_Model_C.csv")


###################################

# Order results based on NLL values
ordered.final.data=order(final.data[,6])
for(j in 1:10) print(final.data[ordered.final.data[j],c(1,3:5)])
```

```
## Iterations      log.p      log.a      log.b 
##     407.00       0.94       0.28       1.00 
## Iterations      log.p      log.a      log.b 
##     502.00       0.96       0.10       0.81 
## Iterations      log.p      log.a      log.b 
##     452.00       0.82       0.29       0.88 
## Iterations      log.p      log.a      log.b 
##     261.00       0.72       0.11       0.53 
## Iterations      log.p      log.a      log.b 
##     287.00       0.74       0.09       0.53 
## Iterations      log.p      log.a      log.b 
##     428.00       0.57       0.10       0.33 
## Iterations      log.p      log.a      log.b 
##     255.00       0.78       0.06       0.54 
## Iterations      log.p      log.a      log.b 
##     333.00       0.99       0.06       0.78 
## Iterations      log.p      log.a      log.b 
##     203.00       0.54       0.06       0.22 
## Iterations      log.p      log.a      log.b 
##     358.00       0.48       0.15       0.24
```

```
####################################

# Run the code for a high number of iterations to make the NLL value more precise.

simulations.check = 1e6
p = 10^final.data[ordered.final.data[1],3]
a = 10^final.data[ordered.final.data[1],4]
b = 10^final.data[ordered.final.data[1],5]
mort <- array(0:0, dim=c(nrow=0,ncol=5))
    
for (i in 1:num.doses) {
    
    something = rbeta(n = simulations.check, shape1 = a, shape2 = b)*p*doses[i]
    lambda.a = something*inoculum.freq
    big.a = which(lambda.a > lambda.limit)
    lambda.a[big.a] = lambda.limit  
    lambda.b = something*(1-inoculum.freq)
    big.b = which(lambda.b > lambda.limit)
    lambda.b[big.b] = lambda.limit              
    infectors.a = rpois(n = simulations.check, lambda = lambda.a)
    infectors.b = rpois(n = simulations.check, lambda = lambda.b)
    infectors.a.1 = rbinom(n = simulations.check, size = (seg.per.virpar*infectors.a), prob = gf.a)
    infectors.a.2 = (seg.per.virpar*infectors.a) - infectors.a.1
    infectors.b.1 = rbinom(n = simulations.check, size = (seg.per.virpar*infectors.b), prob = gf.b)
    infectors.b.2 = (seg.per.virpar*infectors.b) - infectors.b.1
    infectors.1 = infectors.a.1 + infectors.b.1                         
    infectors.2 = infectors.a.2 + infectors.b.2                         
    infected = which(infectors.1 > 0 & infectors.2)             
    survival = (simulations.check - length(infected))/simulations.check                 
    infection.ab = length(which(infectors.a.2[infected] > 0 & infectors.b.2[infected] > 0))/simulations.check
    infection.a = length(which(infectors.a.2[infected] > 0 & infectors.b.2[infected] == 0))/simulations.check   
    infection.b = length(which(infectors.a.2[infected] == 0 & infectors.b.2[infected] > 0))/simulations.check   

    if(survival < 0) survival = 0       
    if(infection.a < 0) infection.a = 0 
    if(infection.b < 0) infection.b = 0
    if(infection.ab < 0) infection.ab = 0
            
    data.for.comp <- c(s.uninfected[i], s.infected.a[i], s.infected.b[i], s.infected.ab[i])
    model.for.comp <- c(survival, infection.a, infection.b, infection.ab)
    likelihood = dmultinom(x = data.for.comp, size = NULL, prob = model.for.comp, log = TRUE)
            
    mort=rbind(mort, c(survival, infection.a, infection.b, infection.ab, likelihood))

}   # end i loop

# NLL value
nll.model.2 = -sum(mort[,5])
print(nll.model.2)
```

```
## [1] 2526.779
```

```
####################################

# Plot the data first
par(mar = c(5, 6, 4, 2))
plot(x = log2(moi.vals), y = f.inf, cex = 1.5,
     main = "Model C", cex.main = 2, font.main = 1,
     xlab = "", ylab = "", 
     type = "p", col = "blue", pch = 19, 
     xlim = c(-4, 4), ylim = c(0, 1),
     cex.lab = 2, cex.axis = 2,
     las = 1)
points(x = log2(moi.vals), y = f.coinf, type = "p", col = "magenta", pch = 19, cex = 1.5)
title(xlab = expression("log"[2]*" MOI"), cex.lab = 2)
title(ylab = expression("Infected and co-infected cells"), line = 3.7, cex.lab = 2)

# Set model parameters
p = 10^final.data[ordered.final.data[1],3]
a = 10^final.data[ordered.final.data[1],4]
b = 10^final.data[ordered.final.data[1],5]

# make array for results, 
mort <- array(0:0, dim=c(nrow=0,ncol=4))

for (i in 1:num.doses.check) {
    something = rbeta(n = simulations.check, shape1 = a, shape2 = b)*p*doses.check[i]
    lambda.a = something*inoculum.freq  # Here the code is modified to limit size of lambda values.
    big.a = which(lambda.a > lambda.limit)
    lambda.a[big.a] = lambda.limit  
    lambda.b = something*(1-inoculum.freq)
    big.b = which(lambda.b > lambda.limit)
    lambda.b[big.b] = lambda.limit              
    infectors.a = rpois(n = simulations.check, lambda = lambda.a)
    infectors.b = rpois(n = simulations.check, lambda = lambda.b)
    infectors.a.1 = rbinom(n = simulations.check, size = (seg.per.virpar*infectors.a), prob = gf.a)
    infectors.a.2 = (seg.per.virpar*infectors.a) - infectors.a.1
    infectors.b.1 = rbinom(n = simulations.check, size = (seg.per.virpar*infectors.b), prob = gf.b)
    infectors.b.2 = (seg.per.virpar*infectors.b) - infectors.b.1
    infectors.1 = infectors.a.1 + infectors.b.1
    infectors.2 = infectors.a.2 + infectors.b.2
    infected = which(infectors.1 > 0 & infectors.2)
    survival = (simulations.check - length(infected))/simulations.check
    infection.ab = length(which(infectors.a.2[infected] > 0 & infectors.b.2[infected] > 0))/simulations.check
    infection.a = length(which(infectors.a.2[infected] > 0 & infectors.b.2[infected] == 0))/simulations.check
    infection.b = length(which(infectors.a.2[infected] == 0 & infectors.b.2[infected] > 0))/simulations.check
  mort=rbind(mort, c(survival, infection.a, infection.b, infection.ab))
        
}   # end i loop
    
# Plot model for this parameter set, infection first
lines(x = log2(doses.check), y = 1-(mort[,1]), col = "blue", lty = 2, lwd = 3)
# Now plot mortality
lines(x = log2(doses.check), y = mort[,4], col = "magenta", lwd = 3)
```

Fitting this model results in marked improvement over Models A and B, as demonstrated by the much lower NLL value and the improved fit upon visual inspection of the data and fitted model. This result suggests that the assumption of heterogeneous cell susceptibility to RVFV is supported by the data.

Finally, to support the graphing of these relationships for the manuscript in any environment, we export the predicted relationship between MOI, infection and co-infection as a \*.csv file.

```
# Aggregate all the results into a single array
model2.results <- cbind(doses.check, mort)
colnames(model2.results) <- c("MOI","Uninfected", "Only eGFP", "Only mCherry", "Coinfected")

# Export the data
setwd("C:/R_Output_EBM/Manuscript (Incomplete RVFV particles)")
write.csv(model2.results, file = "Final_results.csv")
```

In all the plots presented here, the fraction of infected cells experimental data are indicated with blue circles, and the model predictions with the dashed blue line. The fraction of co-infected cells experimental data are indicated with magenta circles, and the model predictions with the solid magenta line.

Of note, results derived from **Model C** were chosen for plotting and visualization of **Fig. 2g** of the manuscript.
