## Supplementary File 2 for "Incomplete bunyavirus particles contribute to within-host spread and between-host transmission": Supplementary File 2.html

Modeling virus spread and the relationship between MOI and co-infection


### Modeling virus spread and the relationship between MOI and co-infection

###### Author: Mark P. Zwart

###### Date: 14 February 2022

##### Supplementary File 2

###### Manuscript: “Incomplete bunyavirus particles contribute to within-host spread and between-host transmission”

###### Erick Bermúdez-Méndez, Kirsten F. Bronsvoort, Mark P. Zwart, Sandra van de Water, Ingrid Cárdenas-Rey, Rianka P. M. Vloet, Constantianus J. M. Koenraadt, Gorben P. Pijlman, Jeroen Kortekaas, Paul J. Wichgers Schreur

###### Intro

Based on information on the packaging of Rift Valley Fever virus (RVFV) genome segments into virus particles, and assumptions on genome complementation between incomplete virus particles, here we make predictions of the infection dynamics of RVFV. As a first step, we want to compare virus populations derived from mammalian and insect cells, for which there are differences in the packaging of genome segments.

###### Function for predicting infection dynamics

We specify a function for letting a population composed of virus particles replicate and spread in a population of cells. Importantly, the simulation is setup such that the composition of virus particles can vary in terms of the segments present in them. This function generates a 3-dimensional array, with the rows being individual sims, columns representing time points, and in the z-dimension 1 is the fraction of infected cells, 2 is the realized MOI, and 3 is the fraction of cells that is infected due to co-infection (i.e. infected cells in which infectious three-segmented virus particles did not contribute to infection). Note that in this version of the simulations, virus populations can go extinct and then need to be removed from subsequent analyses.

```
sim.virus <- function(tot.sims, max.time, ini.inf, vir.prod, inf.prob, num.cells, vp.types) {
  
  # Array to store the data on the number of infected cells, MOI and the 
  # contribution of incomplete particles to infection.
  sim.data <- array(NA, dim = c(tot.sims, max.time, 3))

  # Loop for the simulations
  for(i in 1:tot.sims) {
    
      # Set the number of infected cells to the starting value, exit.flag to zero,
    # and send data for t = 1 to array.
      inf.prev = 0
      inf.now  = ini.inf
      exit.flag = 0
      time = 1
      sim.data[i,time,c(1:3)] = c(ini.inf, NA, NA)
            
      # While loop for growth of the infection, which stops when all cells are 
      # infected or the max time is reached.
      while(exit.flag == 0) {
        
          # Time
          time = time + 1
          
          # Draw the total number of virus particles invading each cell. First 
          # determine the mean number of invading virus particles. Set to zero once 
          # all cells are infected.
          mean.lambda = (inf.now*vir.prod*inv.prob)/(num.cells - inf.now - inf.prev)
          
        # Then draw the number of invading particles for each cell
        lambda.tot = rpois(n = (num.cells - inf.now), lambda = mean.lambda)
          
        # Determine which cells have > 0 vp, so that you only work with these
        invaded = which(lambda.tot > 0)
          lambda.tot.inv = lambda.tot[invaded]
        num.invaded = length(lambda.tot.inv)
        
          # Nested loop drawing the virus particle types for each cell, and 
        # determining whether cells are infected or not.
        
          inf.new = 0
        comp.now = 0    
      
        # Nest this loop which determines the identities of infecting virus
        # particles, so that if lambda is zero the script does not crash (because 
        # size = 0  in sample(), is not allowed), but instead exits the while loop
        # as the population has gone extinct under assumptions made (1 time point
        # window for infecting new cells).
        if(num.invaded > 0) {
          for(j in 1:num.invaded) {
                    
              # Now draw the number of hits for each type of virus particle for this 
              # cell, but only return unique values since we don't care how many of 
              # each type of virus particle is present, only if it is present.
                all.lambdas = unique(sample(x = 1:8, size = lambda.tot.inv[j], 
                                   replace = TRUE, prob = vp.types[,4]))
                
                # Now determine if the cell is infected, and for infected cells 
            # determine whether complementation between virus particles with 
                # an incomplete  set of genome segments was necessary for infection.
                inv.now = rbind(vp.types[all.lambdas,])
                if(sum(inv.now[,1]) > 0 & sum(inv.now[,2]) > 0 & sum(inv.now[,3]) > 0) {
                    inf.new = inf.new + 1
                  if(length(which(all.lambdas == 8)) == 0) comp.now = comp.now + 1
              }
        
            } # end of j loop for this cell
        } else exit.flag = 1 # end of if statement  
                
        # Update numbers
        inf.prev = inf.prev + inf.now
        inf.now = inf.new
        tot.inf = inf.prev + inf.new
        
          # Send data to arrays
        sim.data[i,time,c(1:3)] = c(tot.inf, mean.lambda, (comp.now/inf.new))
          
        # Evaluate condition for while loop
        if(tot.inf >= num.cells) exit.flag = 1
        if(time >= max.time) exit.flag = 1
                
    } # end of while loop
    
  } # end of i loop
  return(sim.data)
}
```

###### General conditions for simulations

We start by defining the general conditions for which we will run the passaging. These parameters have been chosen to be similar to the cell culture conditions, and at the same time, representative for a situation in which a virus population is expanding locally (i.e. the total number of cells is kept small, so that the MOI increases locally as the infection progresses). This situation is representative of a virus replication and expansion in a mass of cells where the movement of virus particles is constrained, as could occur in real life tissues or for example in a plaque assay *in vitro*.

```
# General parameters for the simulations that need to be set.
tot.sims  <- 50     # The total number of simulations to be run for each scenario
max.time  <- 10     # Housekeeping: a maximum number of time steps allowed.

# The biological parameters that we will keep constant over all simulations here.
ini.inf   <- 1      # The number of cells initially infected.
vir.prod  <- 10^3   # The number of virus particles produced per infected cell.
num.cells <- 10^2   # The total number of cells. For exploring the relationship between 
                  # MOI and incomplete particle co-infections, set to a higher 
                  # value (10^3 instead of 10^2).
inv.prob  <- 10^-1.4    # The probability that each virus particle "invades" a new
                      # cell (i.e. that it enters the cell and can contribute 
                      # to infection if all genome segments are present).

# Finally, set random seed for reproducibility.
set.seed(42)
```

###### Predictions in mammalian cells

Here we make predictions for the mammalian cells.

###### Random packager with co-infection by incomplete virus particles

Next, we need to specify what the virus particles look like with respect to the packaging of viral genome segments into virus particles. Based on empirical data we have a good idea of what this distribution looks like for the real virus, so we generate a matrix containing this information. Note that this matrix will also determine infectivity, so it also has bearing on whether incomplete virus particles are infectious (i.e. for simplicity, the generation of incomplete virus particles is set to zero for generating a prediction in which incomplete particles do not contribute to infection spread). First we consider the situation in which we have a random packager in mammalian cells, and that incomplete virus particles can complement each other and thereby cause cellular infection.

```
# The first three columns indicate the presence or absence of a segment, and the
# final column represents the frequency at which that combination is present.
vp.types = array(dim = c(8, 4), data = c(
    0, 1, 0, 0, 1, 0, 1, 1,
    0, 0, 1, 0, 1, 1, 0, 1,
    0, 0, 0, 1, 0, 1, 1, 1,
    0.5, rep(0.09, 3), rep(0.06, 3), 0.05))
colnames(vp.types) <- c("Segment 1", "Segment 2", "Segment 3", "Frequency of VP type")

print(vp.types)
```

```
##      Segment 1 Segment 2 Segment 3 Frequency of VP type
## [1,]         0         0         0                 0.50
## [2,]         1         0         0                 0.09
## [3,]         0         1         0                 0.09
## [4,]         0         0         1                 0.09
## [5,]         1         1         0                 0.06
## [6,]         0         1         1                 0.06
## [7,]         1         0         1                 0.06
## [8,]         1         1         1                 0.05
```

```
# Rename the vp.types array for the random packager, to make it available later 
# on for normalization of the number of virus particles for the selective
# packager.
vp.types.rp = vp.types
```

Now we run the simulations for these conditions, and then first make a plot of the mean, and then of some individual replicates.

```
# Run the simulations using the simulation function
sim.data <- sim.virus(tot.sims = tot.sims, max.time = max.time, ini.inf = ini.inf, 
          vir.prod = vir.prod, inf.prob = inf.prob, num.cells = num.cells,
          vp.types = vp.types)

# Determine in how many cases all cells became infected, and exclude those cases
# where this didn't occur.
keep.these = rep(NA,tot.sims)
for(i in 1:tot.sims) keep.these[i] = max(sim.data[i,,1], na.rm=TRUE) >= num.cells
print(sum(keep.these)) # Report the number of cases in which the population went
```

```
## [1] 41
```

```
                       # extinct. These numbers should be taken into account
                       # when comparing different scenarios.
t.sim.data <- sim.data[keep.these,,]

# Now determine the mean number of infected cells, MOI and fraction of cells 
# infected by incomplete virus particles.
mean.inf = rep(NA, max.time)
mean.moi = mean.inf
mean.coi = mean.inf

for(i in 1:max.time) {
    use.these = which(sim.data[,i,1] > 0)
    mean.inf[i] = mean(sim.data[use.these,i,1])
    mean.moi[i] = mean(sim.data[use.these,i,2])
    mean.coi[i] = mean(sim.data[use.these,i,3])
}

# Rename key outputs so they are still available if the same code is used to 
# explore other conditions further on.
t.sim.data.1 = t.sim.data
mean.inf.1 = mean.inf
mean.moi.1 = mean.moi
mean.coi.1 = mean.coi
```

###### Random packager without co-infection by incomplete virus particles

Next, we consider a random packager in mammalian cells, where incomplete virus particles cannot complement each other and therefore do not cause cellular infection. To this end, we modify the matrix for virus particle composition, setting the frequency of all incomplete variants to zero, and instead replacing them with empty particles.

```
# The first three columns indicate the presence or absence of a segment, and the
# final column represents the frequency at which that combination is present.
vp.types = array(dim = c(8, 4), data = c(
    0, 1, 0, 0, 1, 0, 1, 1,
    0, 0, 1, 0, 1, 1, 0, 1,
    0, 0, 0, 1, 0, 1, 1, 1,
    0.95, rep(0, 6), 0.05))
colnames(vp.types) <- c("Segment 1", "Segment 2", "Segment 3", "Frequency of VP type")

print(vp.types)
```

```
##      Segment 1 Segment 2 Segment 3 Frequency of VP type
## [1,]         0         0         0                 0.95
## [2,]         1         0         0                 0.00
## [3,]         0         1         0                 0.00
## [4,]         0         0         1                 0.00
## [5,]         1         1         0                 0.00
## [6,]         0         1         1                 0.00
## [7,]         1         0         1                 0.00
## [8,]         1         1         1                 0.05
```

Now we run the simulations for these conditions, and then first make a plot of the mean, and then of some individual replicates.

```
# Run the simulations using the simulation function
sim.data <- sim.virus(tot.sims = tot.sims, max.time = max.time, ini.inf = ini.inf, 
          vir.prod = vir.prod, inf.prob = inf.prob, num.cells = num.cells,
          vp.types = vp.types)

# Determine in how many cases all cells became infected, and exclude those cases
# where this didn't occur.
keep.these = rep(NA,tot.sims)
for(i in 1:tot.sims) keep.these[i] = max(sim.data[i,,1], na.rm=TRUE) >= num.cells
print(sum(keep.these)) # Report the number of cases in which the population went
```

```
## [1] 41
```

```
                       # extinct. These numbers should be taken into account
                       # when comparing different scenarios.
t.sim.data <- sim.data[keep.these,,]

# Now determine the mean number of infected cells, MOI and fraction of cells 
# infected by incomplete virus particles.
mean.inf = rep(NA, max.time)
mean.moi = mean.inf
mean.coi = mean.inf

for(i in 1:max.time) {
    use.these = which(sim.data[,i,1] > 0)
    mean.inf[i] = mean(sim.data[use.these,i,1])
    mean.moi[i] = mean(sim.data[use.these,i,2])
    mean.coi[i] = mean(sim.data[use.these,i,3])
}

# Rename key outputs so they are still available if the same code is used to 
# explore other conditions further on.
t.sim.data.2 = t.sim.data
mean.inf.2 = mean.inf
mean.moi.2 = mean.moi
mean.coi.2 = mean.coi
```

###### Selective packager

Finally, we consider a virus that perfectly packages all of its segments into each virus particle produced. To do so, we only need to specify that all segments are present in each virus particle in the matrix for genome segment distributions. However, that would not lead to a fair comparison: to produce the same number of complete virus particles, the perfect packager can require considerably more genome segments available for packaging compared to the random packager. Therefore, we need to determine the number of genome segments available to the random packager and then limit the total number of genome segments available to the selective packager. For simplicity, the total number of virus particles will be kept the same, and a fraction of empty virus particles will be introduced. Note that this means that for this virus, a sensible calculation of the MOI cannot be made with this code.

```
# Determine the total number of genome segments packaged into virus particles
# for the random packager.
vp.segs = rep(NA, 8)
for(i in 1:8) vp.segs[i] = sum(vp.types.rp[i,(1:3)])*vp.types.rp[i,4]

# For a selective packager, the same calculation would render a value of 3x1= 3 
# (3 segments times a frequency of 1), leading to the normalization factor:
norm.pp = sum(vp.segs)/3

# The first three columns indicate the presence or absence of a segment, and the
# final column represents the frequency at which that combination is present.
# Here we consider the effect of a limit to the number of genome segments 
# available, based on the random packager.
vp.types = array(dim = c(8, 4), data = c(
    0, 1, 0, 0, 1, 0, 1, 1,
    0, 0, 1, 0, 1, 1, 0, 1,
    0, 0, 0, 1, 0, 1, 1, 1,
    (1-norm.pp), rep(0, 6), norm.pp))
colnames(vp.types) <- c("Segment 1", "Segment 2", "Segment 3", "Frequency of VP type")
print(vp.types)
```

```
##      Segment 1 Segment 2 Segment 3 Frequency of VP type
## [1,]         0         0         0                 0.74
## [2,]         1         0         0                 0.00
## [3,]         0         1         0                 0.00
## [4,]         0         0         1                 0.00
## [5,]         1         1         0                 0.00
## [6,]         0         1         1                 0.00
## [7,]         1         0         1                 0.00
## [8,]         1         1         1                 0.26
```

Now we run the simulations for these conditions, and then first make a plot of the mean, and then of some individual replicates.

```
# Run the simulations using the simulation function
sim.data <- sim.virus(tot.sims = tot.sims, max.time = max.time, ini.inf = ini.inf, 
          vir.prod = vir.prod, inf.prob = inf.prob, num.cells = num.cells,
          vp.types = vp.types)

# Determine in how many cases all cells became infected, and exclude those cases
# where this didn't occur.
keep.these = rep(NA,tot.sims)
for(i in 1:tot.sims) keep.these[i] = max(sim.data[i,,1], na.rm=TRUE) >= num.cells
print(sum(keep.these)) # Report the number of cases in which the population went
```

```
## [1] 50
```

```
                       # extinct. These numbers should be taken into account
                       # when comparing different scenarios.
t.sim.data <- sim.data[keep.these,,]

# Now determine the mean number of infected cells, MOI and fraction of cells 
# infected by incomplete virus particles.
mean.inf = rep(NA, max.time)
mean.moi = mean.inf
mean.coi = mean.inf

for(i in 1:max.time) {
    use.these = which(sim.data[,i,1] > 0)
    mean.inf[i] = mean(sim.data[use.these,i,1])
    mean.moi[i] = mean(sim.data[use.these,i,2])
    mean.coi[i] = mean(sim.data[use.these,i,3])
}

# Rename key outputs so they are still available if the same code is used to 
# explore other conditions further on.
t.sim.data.3 = t.sim.data
mean.inf.3 = mean.inf
mean.moi.3 = mean.moi
mean.coi.3 = mean.coi
```

###### Plotting the data

We create a single empty plot to represent all the results for the mammalian cells, for all three packaging and infection scenarios.

```
# Create an empty plot for showing all data
par(mar = c(5, 6, 4, 2))

plot(x = c(1, max.time), y = c(0, log10(num.cells)), type = "n",
     main = "Mammalian cells", cex.main = 2, font.main = 1,
     xlab = "Time (viral generations)", ylab = "",
     xlim = c(0,10), ylim = c(0, log10(130)), 
     cex.lab = 2, cex.axis = 2,
     las = 1)
title(ylab = expression("log"[10]*" Infected cells"), line = 3.7, cex.lab = 2)

# Plot individual replicates
num.row = nrow(t.sim.data.1[,,1])
for (i in 1:num.row) lines(x = (0:(max.time-1)), y = log10(t.sim.data.1[i,,1]), 
                           lwd = 0.25, lty = 3, col = "plum")
num.row = nrow(t.sim.data.2[,,1])
for (i in 1:num.row) lines(x = (0:(max.time-1)), y = log10(t.sim.data.2[i,,1]), 
                           lwd = 0.25, lty = 3, col = "lightgreen")
num.row = nrow(t.sim.data.3[,,1])
for (i in 1:num.row) lines(x = (0:(max.time-1)), y = log10(t.sim.data.3[i,,1]), 
                           lwd = 0.25, lty = 3, col = "lightblue")

# Plot the means
lines(x = (0:(max.time-1)), y = log10(mean.inf.1), lwd = 4, lty = 1, 
      col = "purple4")
lines(x = (0:(max.time-1)), y = log10(mean.inf.2), lwd = 4, lty = 1, 
      col = "darkgreen")
lines(x = (0:(max.time-1)), y = log10(mean.inf.3), lwd = 4, lty = 1, 
      col = "darkblue")
```

In this plot, the solid lines represent the mean of simulations for one condition, and the light dotted lines represent individual simulations. The purple lines represent the random packager when co-infection by incomplete particles is allowed, green lines represent the random packager without co-infection by incomplete particles, and the blue lines represent the selective packager.

###### Predictions in insect cells

Next, we make predictions for the insect cell line. We reproduce the script from above, but alter the distribution of genome segments across virus particles, again based on empirical data.

###### Random packager with co-infection by incomplete virus particles

First, we consider the situation in which we have a random packager in insect cells, and that incomplete virus particles can complement each other and thereby cause cellular infection.

```
# The first three columns indicate the presence or absence of a segment, and the
# final column represents the frequency at which that combination is present.
vp.types = array(dim = c(8, 4), data = c(
    0, 1, 0, 0, 1, 0, 1, 1,
    0, 0, 1, 0, 1, 1, 0, 1,
    0, 0, 0, 1, 0, 1, 1, 1,
    0.302, rep(0.07, 3), rep(0.096, 3), 0.2))
colnames(vp.types) <- c("Segment 1", "Segment 2", "Segment 3", "Frequency of VP type")

print(vp.types)
```

```
##      Segment 1 Segment 2 Segment 3 Frequency of VP type
## [1,]         0         0         0                0.302
## [2,]         1         0         0                0.070
## [3,]         0         1         0                0.070
## [4,]         0         0         1                0.070
## [5,]         1         1         0                0.096
## [6,]         0         1         1                0.096
## [7,]         1         0         1                0.096
## [8,]         1         1         1                0.200
```

```
# Rename the vp.types array for the random packager, to make it available later 
# on for normalization of the number of virus particles for the selective
# packager.
vp.types.rp = vp.types
```

Now we run the simulations for these conditions, and then first make a plot of the mean, and then of some individual replicates.

```
# Run the simulations using the simulation function
sim.data <- sim.virus(tot.sims = tot.sims, max.time = max.time, ini.inf = ini.inf, 
          vir.prod = vir.prod, inf.prob = inf.prob, num.cells = num.cells,
          vp.types = vp.types)

# Determine in how many cases all cells became infected, and exclude those cases
# where this didn't occur.
keep.these = rep(NA,tot.sims)
for(i in 1:tot.sims) keep.these[i] = max(sim.data[i,,1], na.rm=TRUE) >= num.cells
print(sum(keep.these)) # Report the number of cases in which the population went
```

```
## [1] 50
```

```
                       # extinct. These numbers should be taken into account
                       # when comparing different scenarios.
t.sim.data <- sim.data[keep.these,,]

# Now determine the mean number of infected cells, MOI and fraction of cells 
# infected by incomplete virus particles.
mean.inf = rep(NA, max.time)
mean.moi = mean.inf
mean.coi = mean.inf

for(i in 1:max.time) {
    use.these = which(sim.data[,i,1] > 0)
    mean.inf[i] = mean(sim.data[use.these,i,1])
    mean.moi[i] = mean(sim.data[use.these,i,2])
    mean.coi[i] = mean(sim.data[use.these,i,3])
}

# Rename key outputs so they are still available if the same code is used to 
# explore other conditions further on.
t.sim.data.4 = t.sim.data
mean.inf.4 = mean.inf
mean.moi.4 = mean.moi
mean.coi.4 = mean.coi
```

###### Random packager without co-infection by incomplete virus particles

Next, we consider a random packager in insect cells, and that incomplete virus particles cannot complement each other and therefore do not cause cellular infection. To this end, we modify the matrix for virus particle composition, setting the frequency of all incomplete variants to zero, and instead replacing them with empty particles.

```
# The first three columns indicate the presence or absence of a segment, and the
# final column represents the frequency at which that combination is present.
vp.types = array(dim = c(8, 4), data = c(
    0, 1, 0, 0, 1, 0, 1, 1,
    0, 0, 1, 0, 1, 1, 0, 1,
    0, 0, 0, 1, 0, 1, 1, 1,
    0.8, rep(0, 6), 0.2))
colnames(vp.types) <- c("Segment 1", "Segment 2", "Segment 3", "Frequency of VP type")

print(vp.types)
```

```
##      Segment 1 Segment 2 Segment 3 Frequency of VP type
## [1,]         0         0         0                  0.8
## [2,]         1         0         0                  0.0
## [3,]         0         1         0                  0.0
## [4,]         0         0         1                  0.0
## [5,]         1         1         0                  0.0
## [6,]         0         1         1                  0.0
## [7,]         1         0         1                  0.0
## [8,]         1         1         1                  0.2
```

Now we run the simulations for these conditions, and then first make a plot of the mean, and then of some individual replicates.

```
# Run the simulations using the simulation function
sim.data <- sim.virus(tot.sims = tot.sims, max.time = max.time, ini.inf = ini.inf, 
          vir.prod = vir.prod, inf.prob = inf.prob, num.cells = num.cells,
          vp.types = vp.types)

# Determine in how many cases all cells became infected, and exclude those cases
# where this didn't occur.
keep.these = rep(NA,tot.sims)
for(i in 1:tot.sims) keep.these[i] = max(sim.data[i,,1], na.rm=TRUE) >= num.cells
print(sum(keep.these)) # Report the number of cases in which the population went
```

```
## [1] 50
```

```
                       # extinct. These numbers should be taken into account
                       # when comparing different scenarios.
t.sim.data <- sim.data[keep.these,,]

# Now determine the mean number of infected cells, MOI and fraction of cells 
# infected by incomplete virus particles.
mean.inf = rep(NA, max.time)
mean.moi = mean.inf
mean.coi = mean.inf

for(i in 1:max.time) {
    use.these = which(sim.data[,i,1] > 0)
    mean.inf[i] = mean(sim.data[use.these,i,1])
    mean.moi[i] = mean(sim.data[use.these,i,2])
    mean.coi[i] = mean(sim.data[use.these,i,3])
}

# Rename key outputs so they are still available if the same code is used to 
# explore other conditions further on.
t.sim.data.5 = t.sim.data
mean.inf.5 = mean.inf
mean.moi.5 = mean.moi
mean.coi.5 = mean.coi
```

###### Selective packager

Finally, we consider a virus that perfectly packages all of its segments into each virus particle produced. To do so, we only need to specify that all segments are present in each virus particle in the matrix for genome segment distributions.

```
# Determine the total number of genome segments packaged into virus particles
# for the random packager.

vp.segs = rep(NA, 8)
for(i in 1:8) vp.segs[i] = sum(vp.types.rp[i,(1:3)])*vp.types.rp[i,4]

# For a perfect packager, the same calculation would render a value of 3x1= 3 
# (3 segments times a frequency of 1), leading to the normalization factor:
norm.pp = sum(vp.segs)/3

# The first three columns indicate the presence or absence of a segment, and the
# final column represents the frequency at which that combination is present.
# Here we consider the effect of a limit to the number of genome segments 
# available, based on the random packager.
vp.types = array(dim = c(8, 4), data = c(
    0, 1, 0, 0, 1, 0, 1, 1,
    0, 0, 1, 0, 1, 1, 0, 1,
    0, 0, 0, 1, 0, 1, 1, 1,
    (1-norm.pp), rep(0, 6), norm.pp))
colnames(vp.types) <- c("Segment 1", "Segment 2", "Segment 3", "Frequency of VP type")
print(vp.types)
```

```
##      Segment 1 Segment 2 Segment 3 Frequency of VP type
## [1,]         0         0         0                0.538
## [2,]         1         0         0                0.000
## [3,]         0         1         0                0.000
## [4,]         0         0         1                0.000
## [5,]         1         1         0                0.000
## [6,]         0         1         1                0.000
## [7,]         1         0         1                0.000
## [8,]         1         1         1                0.462
```

Now we run the simulations for these conditions, and then first make a plot of the mean, and then of some individual replicates.

```
# Run the simulations using the simulation function
sim.data <- sim.virus(tot.sims = tot.sims, max.time = max.time, ini.inf = ini.inf, 
          vir.prod = vir.prod, inf.prob = inf.prob, num.cells = num.cells,
          vp.types = vp.types)

# Determine in how many cases all cells became infected, and exclude those cases
# where this didn't occur.
keep.these = rep(NA,tot.sims)
for(i in 1:tot.sims) keep.these[i] = max(sim.data[i,,1], na.rm=TRUE) >= num.cells
print(sum(keep.these)) # Report the number of cases in which the population went
```

```
## [1] 50
```

```
                       # extinct. These numbers should be taken into account
                       # when comparing different scenarios.
t.sim.data <- sim.data[keep.these,,]

# Now determine the mean number of infected cells, MOI and fraction of cells 
# infected by incomplete virus particles.
mean.inf = rep(NA, max.time)
mean.moi = mean.inf
mean.coi = mean.inf

for(i in 1:max.time) {
    use.these = which(sim.data[,i,1] > 0)
    mean.inf[i] = mean(sim.data[use.these,i,1])
    mean.moi[i] = mean(sim.data[use.these,i,2])
    mean.coi[i] = mean(sim.data[use.these,i,3])
}

# Rename key outputs so they are still available if the same code is used to 
# explore other conditions further on.
t.sim.data.6 = t.sim.data
mean.inf.6 = mean.inf
mean.moi.6 = mean.moi
mean.coi.6 = mean.coi
```

###### Plotting the data

We create a single empty plot to represent all the results for the insect cells, for all three packaging and infection scenarios.

```
# Create an empty plot for showing all data
par(mar = c(5, 6, 4, 2))

plot(x = c(1, max.time), y = c(0, log10(num.cells)), type = "n", 
     main = "Insect cells", cex.main = 2, font.main = 1,
     xlab = "Time (viral generations)", ylab = "",
     xlim = c(0,5), ylim = c(0, log10(130)), 
     cex.lab = 2, cex.axis = 2,
     las = 1)
title(ylab = expression("log"[10]*" Infected cells"), line = 3.7, cex.lab = 2)

# Plot individual replicates
num.row = nrow(t.sim.data.1[,,1])
for (i in 1:num.row) lines(x = (0:(max.time-1)), y = log10(t.sim.data.4[i,,1]), 
                           lwd = 0.25, lty = 3, col = "plum")
num.row = nrow(t.sim.data.2[,,1])
for (i in 1:num.row) lines(x = (0:(max.time-1)), y = log10(t.sim.data.5[i,,1]), 
                           lwd = 0.25, lty = 3, col = "lightgreen")
num.row = nrow(t.sim.data.3[,,1])
for (i in 1:num.row) lines(x = (0:(max.time-1)), y = log10(t.sim.data.6[i,,1]), 
                           lwd = 0.25, lty = 3, col = "lightblue")

# Plot the means
lines(x = (0:(max.time-1)), y = log10(mean.inf.4), lwd = 4, lty = 1, 
      col = "purple4")
lines(x = (0:(max.time-1)), y = log10(mean.inf.5), lwd = 4, lty = 1, 
      col = "darkgreen")
lines(x = (0:(max.time-1)), y = log10(mean.inf.6), lwd = 4, lty = 1, 
      col = "darkblue")
```

In this plot, the solid lines represent the mean of simulations for one condition, and the light dotted lines represent individual simulations. The purple lines represent the random packager when co-infection by incomplete particles is allowed, green lines represent the random packager without co-infection by incomplete particles, and the blue lines represent the selective packager.

###### The relationship between MOI and the importance of co-infection by incomplete virus particles

Finally, we want to consider whether these simulations support a key insight that we have: that MOI will determine how important the role of incomplete particles is for establishing infection. At low MOI, we expect a limited contribution by incomplete particles, because few cells are invaded by more than one virus particle and hence complementation will be rare. At high MOI, all cells will be invaded by a complete, infectious particle, and hence complementation is no longer important for spreading the infection (although incomplete particles will contribute genetically to the spreading virus population). At intermediate MOI, infections caused by incomplete particles are more common, but not all cells have been hit by infectious particles. Hence, we predict that at these intermediate MOIs, incomplete particles play an important role in virus spread.

###### First exploration based on simulations for mammalian and insect cells

To see whether this idea makes sense, we can consider the simulation data we have just generated. Since MOI is not strictly controlled during these simulations, as a first exploration we can simply plot the relationship between realized MOI and the fraction of cells in which infection is caused solely by incomplete particles. We have tracked both of these quantities in the simulations above, and so this plot is easily generated. Of course this analysis only makes sense for the random packager with co-infection scenario, as in all other cases infection by incomplete particles alone is categorically ruled out. We generate a single plot displaying the different results at the end of this section.

We can also try to explore this relationship more systematically. First, we modify the function for simulating the infection process so that it is better suited for our purpose here. The function now only simulates a single round of infection, the MOI is specified as a parameter, and output generated is simplified to just return the mean level of co-infection for the conditions specified. The total number of cells (some of which may be uninfected) is specified, so for low MOIs the number of infected cells will decrease and stochastic effects will increase.

```
sim.virus.2 <- function(tot.sims, moi, num.cells, vp.types) {
  
  # function(tot.sims, max.time, ini.inf, vir.prod, inf.prob, num.cells, vp.types)
  
  # Array to store the data on the number of infected cells, MOI and the 
  # contribution of incomplete particles to infection.
  sim.data <- array(NA, dim = c(tot.sims, 3))

  # Loop for the simulations
  for(i in 1:tot.sims) {
    
        # Then draw the number of invading particles for each cell
    lambda.tot = rpois(n = num.cells, lambda = moi)
      
        # Determine which cells have > 0 vp, so that you only work with these
    invaded = which(lambda.tot > 0)
      lambda.tot.inv = lambda.tot[invaded]
        num.invaded = length(lambda.tot.inv)
    
      # Nested loop drawing the virus particle types for each cell, and 
        # determining whether cells are infected or not.
    
      inf.new = 0
        comp.now = 0    
    
        # Nest this loop which determines the identities of infecting virus
        # particles, so that if lambda is zero the script does not crash (because 
        # size = 0  in sample(), is not allowed), but instead exits the while loop
        # as the population has gone extinct under assumptions made (1 time point
        # window for infecting new cells).
        if(num.invaded > 0) {
      for(j in 1:num.invaded) {
                
              # Now draw the number of hits for each type of virus particle for this 
          # cell, but only return unique values since we don't care how many of 
          # each type of virus particle is present, only if it is present.
            all.lambdas = unique(sample(x = 1:8, size = lambda.tot.inv[j], 
                               replace = TRUE, prob = vp.types[,4]))
            
            # Now determine if the cell is infected, and for infected cells 
            # determined whether complementation between virus particles with 
            # incomplete genomes was necessary for infection.
            inv.now = rbind(vp.types[all.lambdas,])
            if(sum(inv.now[,1]) > 0 & sum(inv.now[,2]) > 0 & sum(inv.now[,3]) > 0) {
                inf.new = inf.new + 1
              if(length(which(all.lambdas == 8)) == 0) comp.now = comp.now + 1
              }
       
        } # end of j loop for this cell
    }   

      # Send data to arrays
    sim.data[i,c(1:3)] = c(inf.new, comp.now, (comp.now/inf.new))
      
    } # end of i loop
  
  func.out = c(mean(sim.data[,3]), sd(sim.data[,3]))
  return(func.out)
}
```

What is neat about the simplified function, is that it shows the relative simplicity of the model. The only “interesting” parameters are MOI and the virus particle description, since the number of simulations and the number of cells just affect how representative and reproducible the result is. Let’s predict and plot the results for the two situations (i.e. different virus particle populations originating from different cell lines) we have considered. First, we set the general parameters to be used for modeling both situations.

```
# Set parameters that will be fixed for both vp.types matrices
tot.sims <- 200           
moi.val <- 10^seq(-1, 3, by = 0.1) # Set a range of MOI values for the sims
num.moi.val <- length(moi.val)
# You can simply fix the total number of cells, but for best results set it
# higher for lower MOIs. Make sure you have the same number of values as for MOI!
# With these settings the script is quite heavy and will need to run for a day
# or so. 
num.cells.val <- c(rep(10000, 11), rep(5000, 10), rep(500, 20))

# Some suggested alternative values for a very quick run
# tot.sims = 20
# num.cells.val <- c(rep(1000,11), rep(20, 30))
```

Now set the virus particle composition again, first for mammalian cells and then for insect cells, and run a loop to generate predictions over this range of MOI values. Finally, we can plot the data from both the original simulations over multiple passages and the single passage results with controlled MOI. For simplicity, here it will only be plotted the single passage results with controlled MOI, but if interested, the code for plotting the original simulations over multiple passages is made available within the below chunk of code as a comment.

```
# The first three columns indicate the presence or absence of a segment, and the
# final column represents the frequency at which that combination is present.
vp.types = array(dim = c(8, 4), data = c(
    0, 1, 0, 0, 1, 0, 1, 1,
    0, 0, 1, 0, 1, 1, 0, 1,
    0, 0, 0, 1, 0, 1, 1, 1,
    0.5, rep(0.09, 3), rep(0.06, 3), 0.05))
colnames(vp.types) <- c("Segment 1", "Segment 2", "Segment 3", "Frequency of VP type")

# Generate an array to store the results
coi.res.1 <- array(NA, dim = c(num.moi.val, 3))
colnames(coi.res.1) <- c("MOI", "Coinfection", "SD_Coinfection")

# Run the loop
for(i in 1:num.moi.val) {
  
  # Set the MOI and num cells
  moi <- moi.val[i]   
  num.cells = num.cells.val[i]
  
  # Run function and send results to array
  coi.res.1[i,] <- c(moi, sim.virus.2(tot.sims = tot.sims, moi = moi, 
                                    num.cells = num.cells, vp.types = vp.types))
}

# Now we repeat the above simulations for virus particles produced in insect 
# cells. The code from above is re-used, renaming the results array so that both 
# results are available for graphing.
vp.types = array(dim = c(8, 4), data = c(
    0, 1, 0, 0, 1, 0, 1, 1,
    0, 0, 1, 0, 1, 1, 0, 1,
    0, 0, 0, 1, 0, 1, 1, 1,
    0.302, rep(0.07, 3), rep(0.096, 3), 0.2))
colnames(vp.types) <- c("Segment 1", "Segment 2", "Segment 3", "Frequency of VP type")

coi.res.2 <- array(NA, dim = c(num.moi.val, 3))
colnames(coi.res.2) <- c("MOI", "Coinfection", "SD_Coinfection")

for(i in 1:num.moi.val) {
  moi <- moi.val[i]   
  num.cells = num.cells.val[i]
  coi.res.2[i,] <- c(moi, sim.virus.2(tot.sims = tot.sims, moi = moi, 
                                    num.cells = num.cells, vp.types = vp.types))
}

# Now let's plot the different results. First, we make an empty plot.
par(mar = c(5, 6, 4, 2))

plot(x = 0, y = 0, type = "n", 
     xlab = expression("log"[10]*" MOI"), ylab = "", 
     xlim = c(-1, 3), ylim = c(0, 1),
     cex.lab = 2, cex.axis = 2,
     las = 1)
title(ylab = "Incomplete particles infection", line = 4, cex.lab = 2)

# Next we add the data from the original simulations (MOI not fixed).
#points(x = log10(t.sim.data.1[,,2]), y = t.sim.data.1[,,3], pch = 1, 
#       col = "orange")
#points(x = log10(t.sim.data.4[,,2]), y = t.sim.data.4[,,3], pch = 2, 
#       col = "red")

# Finally we add the predictions for the controlled MOI simulations.
lines(x = log10(coi.res.1[,1]), y = coi.res.1[,2], lty = 1, lwd = 4, col = "orange")
lines(x = log10(coi.res.2[,1]), y = coi.res.2[,2], lty = 1, lwd = 4, col = "darkred")
```

The lines represent the results for a systematic test over a range of controlled MOI values, with orange corresponding to the mammalian cells and red to the insect cells. There are some clear trends: at low MOI there are few infections caused by co-infection (with a high varation in the individual simulations due to a sampling effect). At high MOI, infections caused by incomplete particles indeed become rare, whereas they are most common at intermediate MOI (i.e. between 1 and 10).

###### A more general approach

These results look quite promising and are congruent with our intuition, and the neat thing about the predictions is that there are no free variables to be estimated. However, the results do depend on the distribution of genome segments over virus particles (i.e. the *vp.types* matrix). As a final check of the generality of these results, we can therefore consider the relationship between MOI and the frequency of infection initiated solely by incomplete particles for a wide distribution of genome segments over virus particle types. The code below draws random values for the abundance of different virus particle types, and then determines the MOI vs. co-infection relationship.

First, we set some general parameters for this exploration. We can re-use the sim.virus.2() function and the parameters set for this function above. Then we have a loop that draws random values for the distribution of virus particle types, evaluates the relationship between MOI and co-infections, and plots the results.

```
# The number of times to randomly draw values for vp.types and evaluate them.
vp.sims = 40

# Generate an array to store the results. First array is for the co-infection 
# results, the second array to record the vp.types drawn.
sim.coinf <- array(NA, dim = c(vp.sims, num.moi.val))
sim.vptyp <- array(NA, dim = c(vp.sims, 8)) 
# Create a template for the vp.types array
# The first three columns indicate the presence or absence of a segment, and the
# final column represents the frequency at which that combination is present.
vp.types.tem = array(dim = c(8, 4), data = c(
    0, 1, 0, 0, 1, 0, 1, 1,
    0, 0, 1, 0, 1, 1, 0, 1,
    0, 0, 0, 1, 0, 1, 1, 1,
    rep(NA, 8)))
colnames(vp.types.tem) <- c("Segment 1", "Segment 2", "Segment 3", "Frequency of VP type")

# Main loop 
for(i in 1:vp.sims) {
  
  # print(i)
  
  # Select random values for vp.types array. We draw a random number from a 
  # uniform distribution for each segment frequency, and then normalize by the 
  # sum of all drawn frequencies.
  vp.types <- vp.types.tem
  rand.val <- runif(n = 8)
  norm.rand.val = rand.val/sum(rand.val) 
  vp.types[,4] <- norm.rand.val
  
  # Now we run the simulations for determining the relationship between MOI and 
  # co-infection. Make an array to store the data and run the loop for 
  # determining co-infection over MOI values.
  coi.res <- array(NA, dim = c(num.moi.val, 3))
  
  for(j in 1:num.moi.val) {
    
    # Set the MOI and num cells
    moi <- moi.val[j]   
    num.cells = num.cells.val[j]
     
    # Run function and send results to array
    coi.res[j,] <- c(moi, sim.virus.2(tot.sims = tot.sims, moi = moi, 
                        num.cells = num.cells, vp.types = vp.types))
  }
  
  # Send results to arrays
  sim.coinf[i,] <- coi.res[,2]
  sim.vptyp[i,] <- norm.rand.val
  
}

# Export the data in case you want to graph it some other way.
setwd("C:/R_Output_EBM/Manuscript (Incomplete RVFV particles)")
write.csv(sim.coinf, file = "Sim_Coinf.csv")
write.csv(sim.vptyp, file = "Sim_Vptype.csv")


# Graph the results. Make an empty plot.
par(mar = c(5, 6, 4, 2))

plot(x = 0, y = 0, type = "n", 
     xlab = expression("log"[10]*" MOI"), ylab = "",
     xlim = c(-1, 3), ylim = c(0, 1), 
     cex.lab = 2, cex.axis = 2,
     las = 1)
title(ylab = "Incomplete particles infection", line = 3.7, cex.lab = 2)

# Then determine the frequency of virus particles with the complete genome, 
# since this will be a predictor of the importance of co-infection. Make colors 
# for displaying the lines according to the content of complete virus particles.
# Note: 2*vp.sims colors made so that only the darker part of the spectrum is
# used to make the lines more clearly visible.
library("RColorBrewer")
order.vp = order(sim.vptyp[,8])
col.palette = colorRampPalette(brewer.pal(9,"YlOrRd"))(2*vp.sims)
lty.val = 1

# Describe the prevalence of complete virus particles for the 40 randomly drawn
# virus particle distributions
min(sim.vptyp[,8]) # minimum frequency of complete particles
```

```
## [1] 0.02282569
```

```
max(sim.vptyp[,8]) # maximum frequency of complete particles
```

```
## [1] 0.288635
```

```
mean((sim.vptyp[,8])) # mean frequency of the complete particles ~ 1/8
```

```
## [1] 0.133986
```

```
# Add the data
for(i in 1:vp.sims) {
  lines(x = log10(moi.val), y = sim.coinf[order.vp[i],], lwd = 3,
        col = col.palette[(vp.sims+i)], lty = lty.val)
}
```

In the figure, the simulation results are ranked by the frequency of complete infectious particles (those containing all three viral genome segments). For the largest fractions the lines are colored in dark shades of red and for the smallest fractions the lines are colored in lighter yellow hues. The resulting figure clearly illustrates that the relationship between MOI and the number of infections caused by incomplete virus particles is quite general, as for each of the conditions simulated there is a bell shaped curve with an optimum around MOI values of 3-10. However, the fraction of complete infectious particles appears to be a determinant of the importance of co-infection by incomplete particles. The different simulation results are therefore consistent with our expectations.
